# Temporally and Spatially Regulated Collagen XVIII Isoforms Impact Ureteric Patterning Through Their TSP1-like Domain

**DOI:** 10.1101/2021.03.08.434365

**Authors:** Mia M. Rinta-Jaskari, Florence Naillat, Heli J. Ruotsalainen, Saad U. Akram, Jarkko T. Koivunen, Valerio Izzi, Veli-Pekka Ronkainen, Takako Sasaki, Ilkka Pietilä, Harri P. Elamaa, Inderjeet Kaur, Aki Manninen, Seppo J. Vainio, Taina A. Pihlajaniemi

**Author notes:** Author for correspondence: Taina Pihlajaniemi, Faculty of Biochemistry and Molecular Medicine, University of Oulu, Aapistie 7, 90230 Oulu, Finland.

## Abstract

Collagen XVIII (ColXVIII) is a component of the extracellular matrix implicated in embryogenesis and control of homeostasis. We provide evidence that ColXVIII has a specific role in kidney ontogenesis by regulating the interaction between mesenchymal and epithelial tissues as observed in analyses of total and isoform-specific knockout embryos, mice, and *ex vivo* organ primordia. ColXVIII deficiency, both temporally and spatially, impacts the 3D pattern of ureteric tree branching morphogenesis *via* its specific isoforms. Proper development of ureteric tree depends on a tight control of the nephron progenitor cells (NPCs). ColXVIII-deficient NPCs are leaving the NPC pool faster than in controls. Moreover, the data suggests that ColXVIII mediates the kidney epithelial tree patterning *via* its N-terminal domains, and especially the Thrombospondin-1-like domain, and that this morphogenetic effect involves ureteric epithelial integrins. Altogether, the results propose a significant role for ColXVIII in a complex signalling network regulating renal progenitors and kidney development.

## INTRODUCTION

The extracellular matrix (ECM) plays a critical role in embryonic development as well as in maintaining organ homeostasis and regulating stem cell fate (Andrew & Ewald, 2010; Lu, Takai, Weaver, & Werb, 2011; Lu, Weaver, & Werb, 2012). The ECM consists of two major compartments, namely the interstitial matrix and the basement membranes (BM) (Andrew & Ewald, 2010; Lu et al., 2011; Lu et al., 2012). Collagen XVIII (ColXVIII) specifically occurs at the BMs and exerts its function *via* three isoforms, and it also serves as a heparan sulphate proteoglycan. The isoforms differ in size and are established on account of two promoters and alternative splicing (Halfter, Dong, Schurer, & Cole, 1998; Muragaki et al., 1995; Rehn & Pihlajaniemi, 1994; Rehn, Hintikka, & Pihlajaniemi, 1996). All isoforms contain a common collagenous portion, a C-terminal domain containing endostatin, and an N-terminal thrombospondin-1 (TSP1)-like domain. The so called medium and long isoforms have an N-terminal mucin-like domain (MUCL-C18) that is flanked in the long form by a Frizzled-like domain which is involved in control of Wnt signalling (Fig. 1) (Kaur et al., 2018; Muragaki et al., 1995; O’Reilly et al., 1997; Saarela, J., Ylikärppä, Rehn, Purmonen, & Pihlajaniemi, 1998).

**Figure 1.**
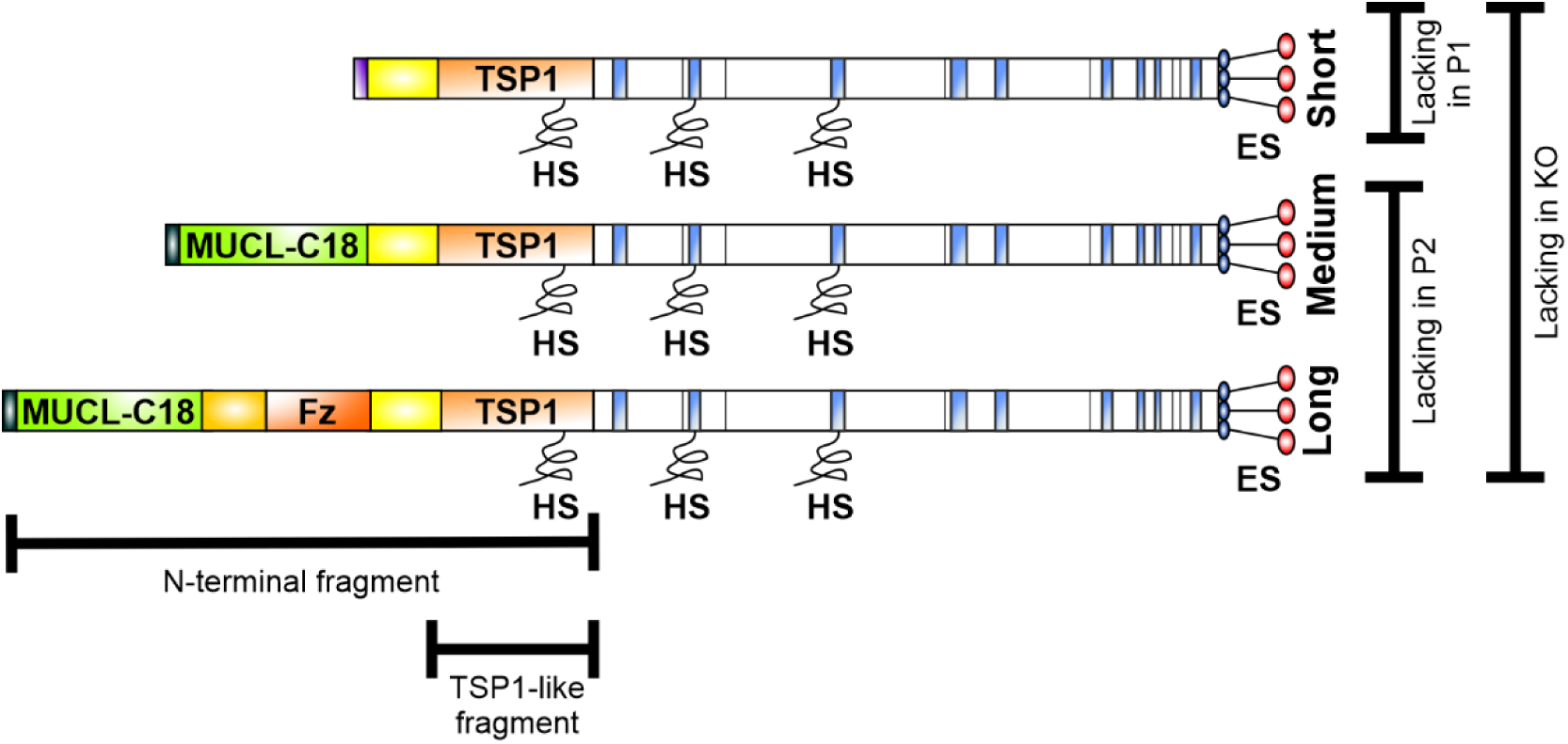
Schematic picture of the structure of ColXVIII and mouse lines used in the study. All three isoforms have a common collagenous domain, C-terminal endostatin domain (ES) and N-terminal TSP1-like domain (TSP1). In addition, the medium and long isoforms have MUCL-C18 domain in the N-terminus, but only the longest isoform has also the N-terminal Frizzled-domain (Fz). The total ColXVIII KO mice lack all three isoforms, whereas the P1 mice lack only the short isoform and the P2 mice lack only the two longer isoforms. The picture also shows the N-terminal fragment and the TSP1-like fragment used in the kidney organ culture studies. HS = heparan sulphate side chain.

In humans, the lack of ColXVIII results in Knobloch syndrome, which is characterised by eye abnormalities and an occipital encephalocele (Caglayan et al., 2014; Passos-Bueno et al., 2006). In addition, other symptoms, including kidney abnormalities, have been described in these patients (Bishop et al., 2010; Passos-Bueno et al., 2006; Seppinen & Pihlajaniemi, 2011; Sertié et al., 1996), in line with the phenotypes noted in the ColXVIII-deficient mice (Aikio et al., 2013; Aikio et al., 2014; Bishop et al., 2010; Kinnunen et al., 2011; Olsen et al., 2002; Utriainen et al., 2004).

In the adult mouse kidney, all three ColXVIII isoforms are expressed, but their localisations differ. The short form is expressed in the tubular BMs, the Bowman’s capsule and the glomerular endothelial BM, whereas the other two localise to the BM of the podocytes, the mesangium and the BM of the kidney vessels (Elamaa, Snellman, Rehn, Autio-Harmainen, & Pihlajaniemi, 2003; Kinnunen et al., 2011; Muragaki et al., 1995; Saarela, J. et al., 1998; Saarela, J., Rehn, Oikarinen, Autio-Harmainen, & Pihlajaniemi, 1998).

With regards of developing mammalian kidney ColXVIII is present in the ureter bud (UB) from E10.5 onwards when the bud goes on to invade the predetermined and committed metanephric mesenchyme (MM) containing the nephron precursor cells (NPCs). ColXVIII has been speculated to have a role in the UB to respond to the mesenchymal signals that control the UB bifurcation during ontogenesis (Karihaloo et al., 2001; Lin et al., 2001).

To target the roles of type XVIII collagen in detail the expression patterns of its three isoforms during renal development were characterised, and sophisticated *ex vivo* and *in vivo* experiments were implemented for better understanding of ColXVIII function in the embryonic kidneys. For these purposes, ColXVIII total and isoform specific knockout embryos and mice (Fig. 1) were used. We now provide evidence that ColXVIII knockout impacts greatly the UB branching process and concurrently reduces the NPC population. This study is the first to describe a role for the non-collagenous N-terminal domain of ColXVIII in renal development where it rescues the noted epithelial malformations in UB. Our study points to a role of the TSP1-like domain of ColXVIII in regulating the UB branching, a process potentially mediated by integrins.

## RESULTS

### Only the short ColXVIII isoform is expressed during the early phase of kidney development, whereas the expression of the two longer forms is detected in more mature structures

While it has been shown that the different ColXVIII isoforms have specific locations in adult mice and human as well as human fetal kidneys (Kinnunen et al., 2011; Saarela, J. et al., 1998), the isoforms’ temporal expression patterns during renal organogenesis have been unknown. Thus, the RNA expression of ColXVIII and its different isoforms was analysed by qPCR from separated MM and UB of the isoform specific ColXVIII knockout embryos lacking either the short isoform (P1) or the two longer isoforms (P2) and ColXVIII total knockout (KO) embryos at E11.5. Two sets of primers were used to ascertain the expression patterns, those directed against the endostatin domain common to all isoforms and those specific to the short isoform. With the endostatin primers, ColXVIII expression was detected both in the MM and the UB of WT and P2 kidneys (Fig. 2a,b), while this expression was lacking in the KO and P1 kidneys (Fig. 2a,b). This expression pattern was confirmed with the short isoform-specific primers (Fig. 2c,d). The results of the qPCR analyses indicate that only the short ColXVIII isoform is expressed in both the MM and UB and that there is no expression of the two longer forms during this early phase of development.

**Figure 2.**
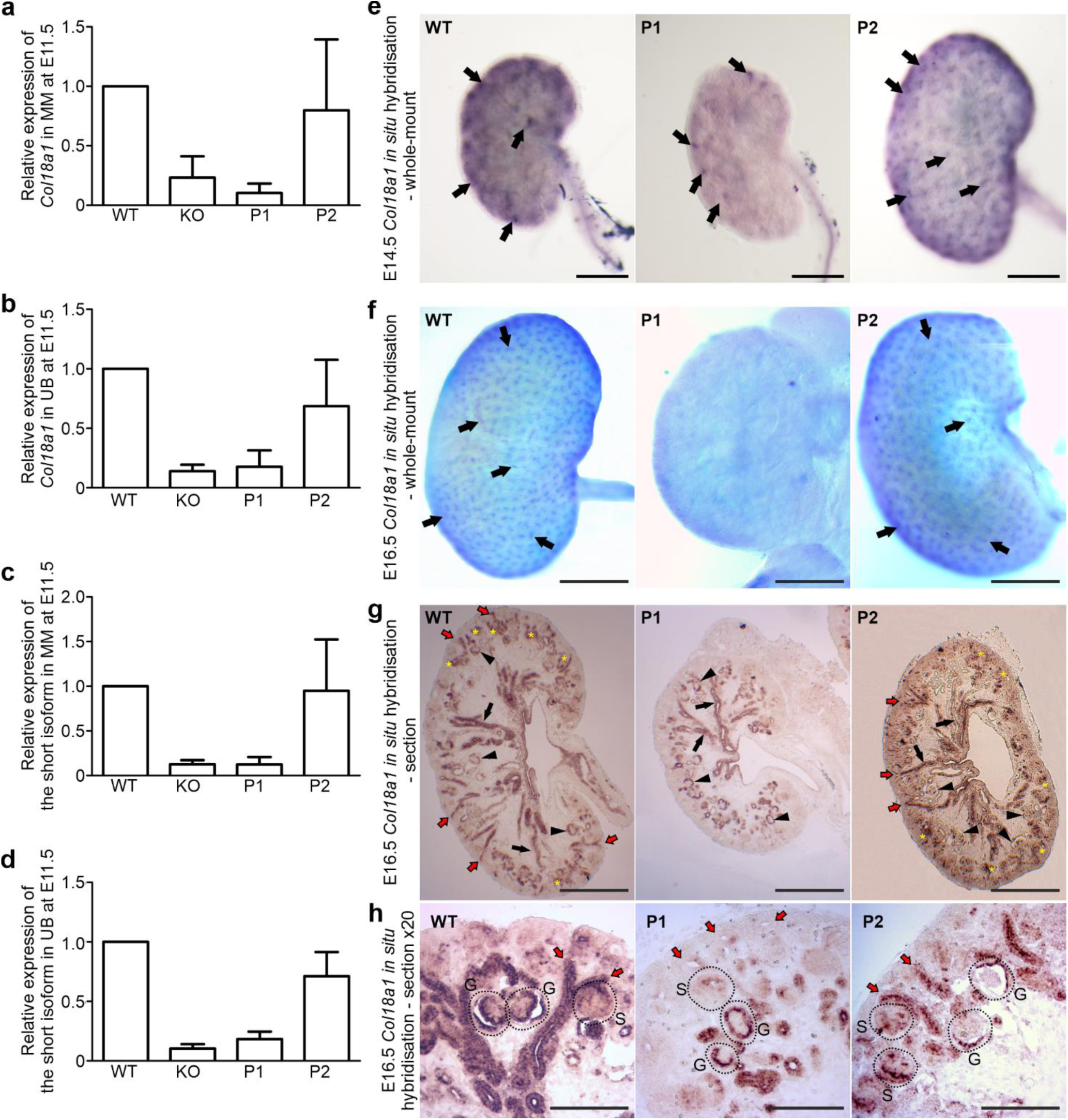
Only the short ColXVIII isoform was expressed during the early stages of kidney development and in the developing structures. The qPCR analyses of separated (a) metanephric mesenchyme (MM) and (b) ureteric bud (UB) at E11.5 using primers from the common endostatin domain of ColXVIII were conducted. The expression of only the short form in the (c) MM and the (d) UB at E11.5 was confirmed with qPCR using specific primer for the short ColXVIII form in KO, P1, and P2; n(WT)=20, n(KO)=18, n(P1)=14, and n(P2)=22 pooled samples. Whole-mount in situ hybridisation of (e) E14.5 (bar: 500 µm) and (f) E16.5 (bar: 500 µm) WT, P1, and P2 kidneys with an antisense ColXVIII RNA probe. Arrows indicate the ColXVIII-positive structures. (g) Section in situ hybridisations of E16.5 WT, P1, and P2 kidneys for ColXVIII expression (bar: 500 µm) and magnifications of the E16.5 section in situ hybridisations (h) (bar: 100 µm). In (g) black arrowheads indicate ColXVIII-positive glomeruli, black arrows indicate collecting ducts, red arrows indicate branching tubules, and yellow stars indicate ColXVIII-positive comma- and S-shape bodies. In (h), red arrows indicate branching tubules, and dashed areas indicate glomeruli (G) and S-shape bodies (S). In graphs, columns indicate mean±s.d.

To visualise the expression of the ColXVIII in the different knockout mice, the whole-mount *in situ* hybridisation of WT and ColXVIII mutant kidneys was carried out that revealed a well-regulated expression of ColXVIII isoforms. At E12.5, the ColXVIII expression was detected in the epithelium of the ureter extending through the kidney in both the WT and the P2 kidneys expressing the short form, while in the P1 kidneys expressing the medium and long forms, the expression was only detectable in the ureteric stalk area (Fig. 2 – figure supplement 1a). At E13.5 (Fig. 2 – figure supplement 1b) and E14.5 (Fig. 2e – figure supplement 1c) the P1 kidneys expressing only the two longer isoforms had a less positive signal in the cortex compared with the WT and P2 kidneys expressing the short isoform. Furthermore, at E16.5, the P1 kidneys expressing only the two longer isoforms were devoid of a ColXVIII expression on the surface of the organ, similarly to the KO kidneys, whereas WT and P2 kidneys expressing the short isoform showed a strong ColXVIII expression in the ureter tips (Fig. 2f).

The section *in situ* hybridisation of E14.5 (Fig. 2 – figure supplement 1d) and E16.5 (Fig. 2g,h) kidneys revealed at both time points positive ColXVIII expression throughout the cortex in the WT and P2 kidneys expressing only the short isoform, including the mature collecting ducts and glomeruli, as well as the glomerular precursors and branching tubules in the nephrogenic zone. In contrast, in the P1 kidneys lacking the short isoform ColXVIII was expressed only in the mature glomeruli and tubules, but the developing structures in the nephrogenic zone were devoid of ColXVIII expression (Fig. 2g,h – figure supplement 1d). ColXVIII immunostaining of E14.5 (Fig. 3) kidney sections confirmed these differences in the isoforms’ expression patterns and clearly indicated that only the short ColXVIII isoform is expressed in the BMs of renal vesicles and comma- and S-shape bodies, as well as in the BMs around the ureteric tips separating ureteric tip cells and NPCs. Moreover, these studies revealed that the isoforms have different localisations in the mature glomeruli: the longer ColXVIII variants were expressed by the podocytes and the short variant localised to the Bowman’s capsule, especially to the layer of parietal epithelial cells, in addition to the glomerular BM (Fig. 3d). In conclusion, the short ColXVIII isoform is the only form expressed in the early phases of renal development, in the ureteric tips and stalks as well as in all stages of the nephron development while the two longer forms are expressed only later in development in the ureteric stalks and in the nephrons which have passed the S-phase.

**Figure 3.**
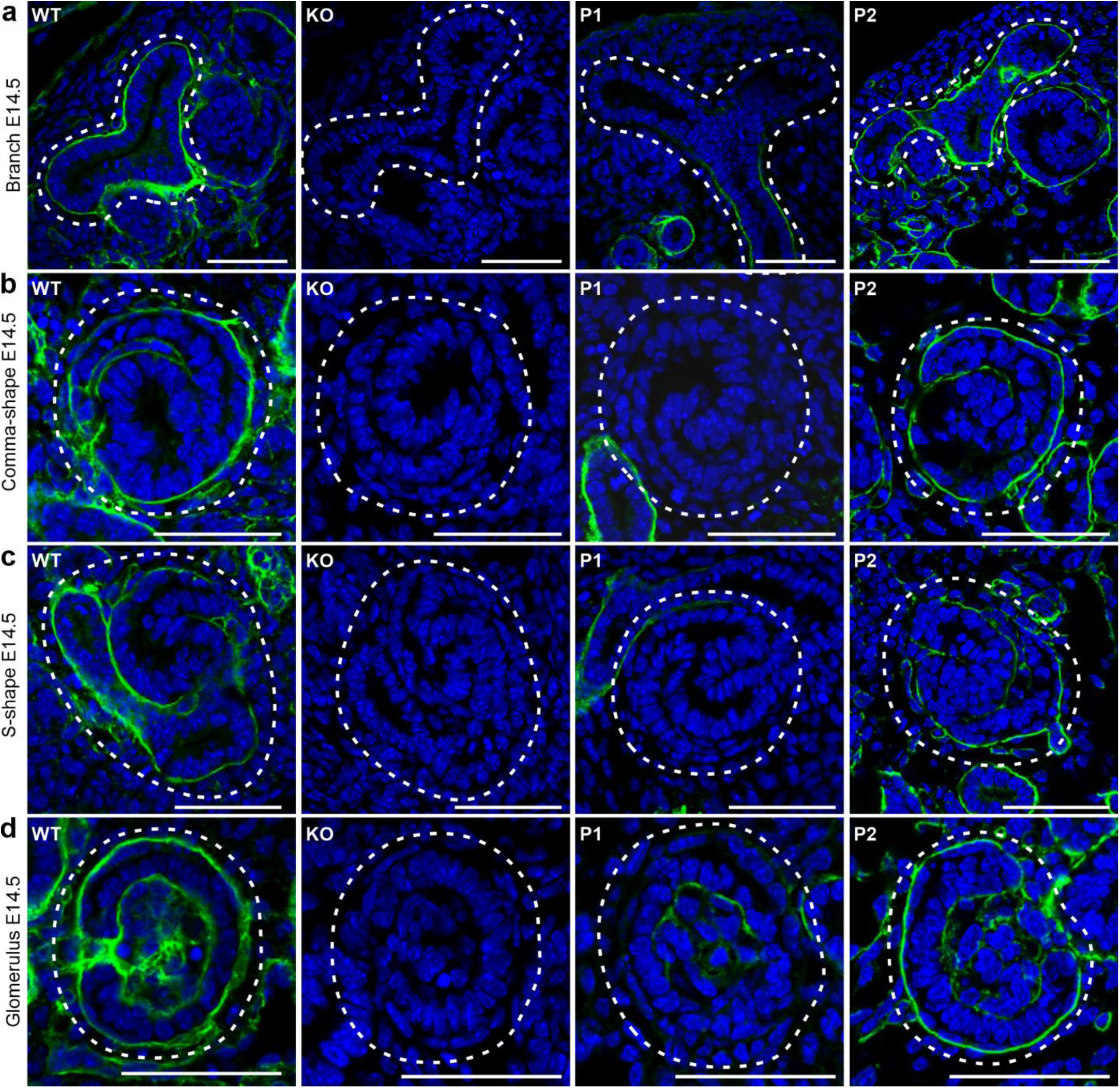
The short form of ColXVIII was the main form expressed in the developing renal structures, and the different ColXVIII isoforms had different localisations in mature glomeruli. Immunostaining for ColXVIII expression in E14.5 kidneys of WT, ColXVIII KO, and P1 and P2 promoter-specific ColXVIII mutant mice. (a) tips of the branching tubules, (b) comma-shape bodies, (c) S-shape bodies of developing glomeruli, and (d) more mature glomeruli. Note that the ColXVIII expression is detected throughout the BM of the ureteric tips in WT and P2 mice expressing the short isoform, where the BM lines both NPCs and ureteric tip cells. Dashed areas indicate the corresponding structures. Bar: 50 μm

### Lack of ColXVIII reduces the number of the NPCs in the nephrogenic niches

ColXVIII is involved in the control of the survival and proliferation of stem cells during adipogenesis in adult mice and such result raised the possibility that similar behavior might be observed in the embryonic kidneys in NPCs (Aikio et al., 2014). We first tested the hypothesis that ColXVIII would have a role in these cells by lineage tracing of Wnt4 positive cells and using live microscopy to investigate NPC behaviour. Wnt4 is one of the earliest known markers of progenitor commitment and most of the NPCs that express Wnt4 commit to nephron formation (Lawlor et al., 2019). The *Wnt4^Cre^*, *Rosa^LacZ^*, and *Rosa^YFP^* transgenic mouse lines were crossed with the *Col18a1^-/-^* (KO) background to follow the behaviour of the Wnt4+ committed NPCs in the ColXVIII-deficient fetuses.

In early kidney development at E11.5 and in the absence of ColXVIII, the Wnt4+ cells were visualized first by β-galactosidase staining. At this time point, the Wnt4+ cells in the *Wnt4^Cre^*; *Rosa^LacZ^*; *Col18a1^+/+^*(WT) embryos were noted to be near the metanephric kidney, but in the absence of ColXVIII LacZ-positive cells localised only to the dorsal portion of the mesonephros/gonad primordia (Fig. 4). ColXVIII deficiency thus delayed the appearance of the Wnt4 signal in the area of the metanephric region at the initiation of renal organogenesis.

**Figure 4.**
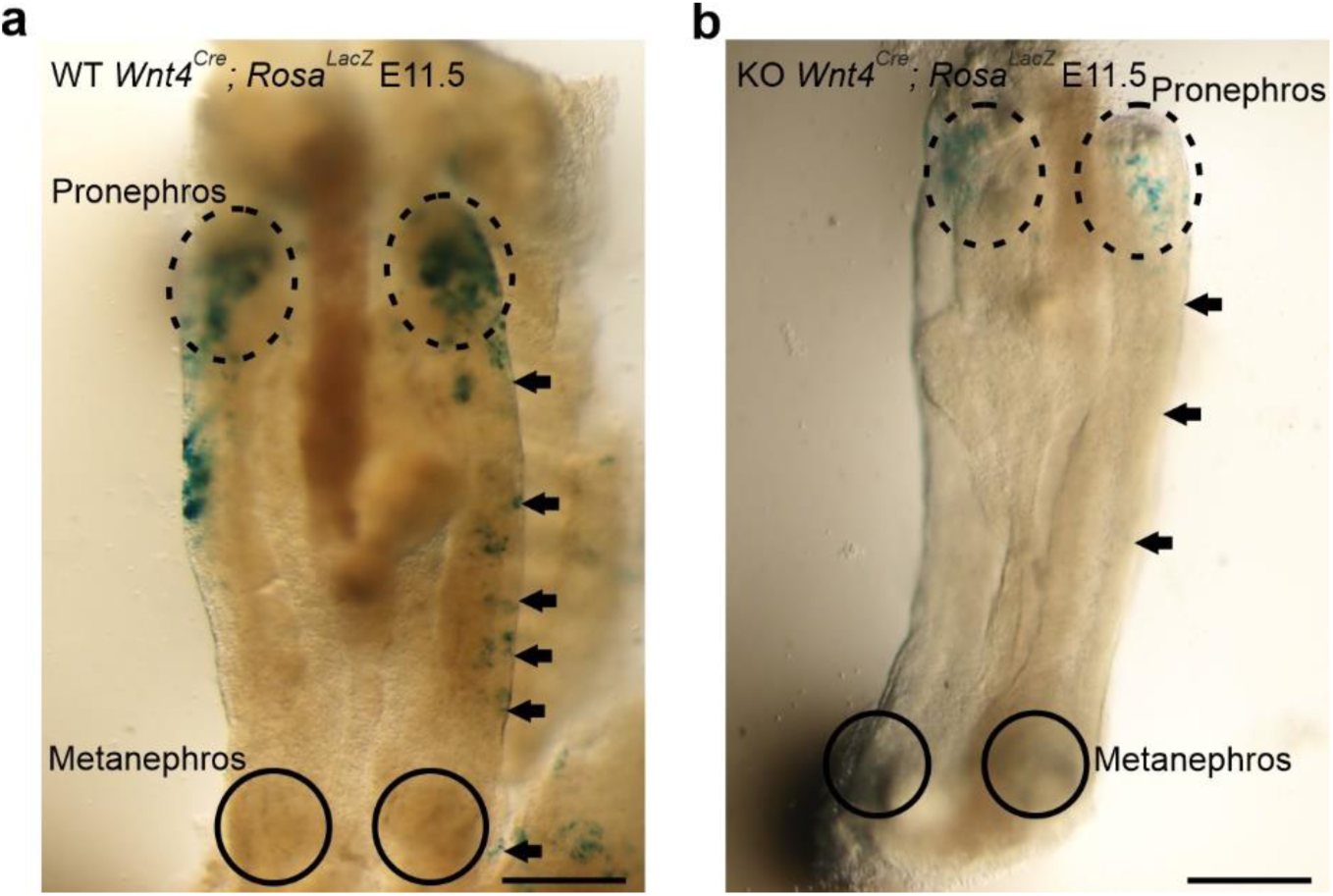
The appearance of Wnt4-positive NPCs was delayed in ColXVIII KO embryos at E11.5. (a) Whole-mount WT AGM of *Wnt4Cre; RosaLacZ* transgenic mice was stained with LacZ (blue color) to visualise Wnt-positive (Wnt4+) cells. The Wnt4+ cells were seen throughout the gonad, including the area next to the forming kidney. (b) Only one Wnt4+ cell was detected in the middle of the mesonephros of E11.5 ColXVIII KO AGM, and no positive cells were seen in the developing kidney area or near it. Arrows indicate Wnt4+ cells localisation on the gonad-mesonephros-metanephros area. Circles indicate the areas where the kidneys are forming. Dashed circles indicate the pronephric area. Bars: 500 μm.

We then prepared the *Rosa^YFP^*; *Wnt4^Cre^*-positive E10.5 aorta-gonad-mesonephros (AGMs) from WT (Fig. 5a, b) and KO (Fig. 5c,d) embryos and set these explants to time-lapse imaging culture for at least 38 hours. In this setting, in the WT embryonic AGMs the Wnt4+ cells were detected in the MM by 5–34 h (mean 24 h), whereas in the KO primordia, such cells were noted by 17–46 h (mean 30.44 h) (Fig. 5e, Movie 1 and 2) suggesting a delay in propagation of the Wnt4+ cells during emergence of the early tubular network.

**Figure 5.**
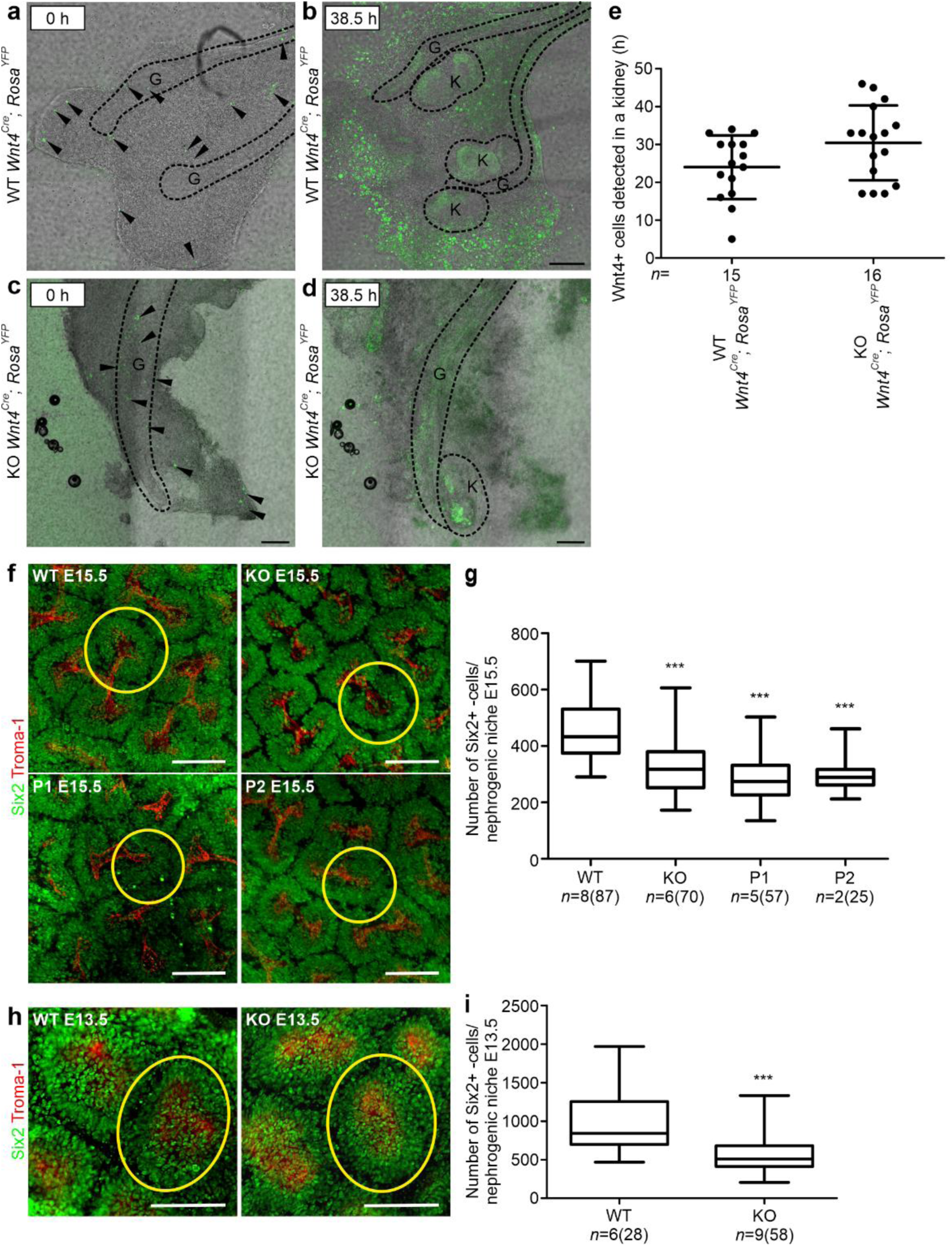
ColXVIII regulated NPC population during kidney development. (a-d) Examples of the aorta-gonad-mesonephros (AGM) cultures of E10.5 *Rosa^YFP^*; *Wnt4^Cre^*-positive WT and KO embryos. In the beginning, the Wnt4+ cells (green, indicated by arrow heads) were seen in the mesonephros/gonad area (outlined, G) and in the limb area in the WT (a) and KO (c). Neither the mesenchyme nor the ureter bud of the kidney had yet formed. The kidneys (K) formed during the culture and became clearly positive for Wnt4 both in the WT (b) and KO (d). Bars: 200 µm. (e) Analyses of the AGM cultures showed that the appearance of the Wnt4+ cells in the kidney was delayed in the ColXVIII-deficient embryos (n=16) when compared with WT embryos (n=15). Positivity was calculated from the moment when the first positive cell was detected in the developing kidney. P-value for the Mann-Whitney U test was 0.0844 (not significant). Multiphoton microscopy maximum intensity projection images of the cortex of Six2 (green)-TROMA-1 (red) whole-mount stained E15.5 (f) and E13.5 (h) kidneys. A yellow circle indicates a typical nephrogenic niche, all of which were separately calculated. Bar: 100 µm. The number of Six2+ NPCs in nephrogenic niches was decreased in the E15.5 (g) ColXVIII mutant mice lacking all forms (KO), the short form (P1), and the medium and long forms (P2), and in the E13.5 (i) KO kidneys. In the E15.5 analyses: n(WT)=8 (87 tips), n(KO)=6 (70 tips), n(P1)=5 (57 tips), and n(P2)=2 (25 tips), and in the E13.5 n(WT)=6 (28 tips) and n(KO)=9 (58 tips). In graphs, the lines in (e) indicate mean±s.d., and in (g) and (i), the boxes show 1^st^ and 3^rd^ quartiles with mean and whiskers indicate min and max values. ***P<0.0001 (Mann-Whitney U -test).

The Wnt4+ cells at E11.5 and E13.5 in the *Rosa^YFP^*; *Wnt4^Cre^*-positive WT and KO kidney sections were stained for yellow fluorescence protein (YFP), Sine oculis-related homeobox 2 (Six2), a marker of NPCs in the cap mesenchyme (CM) (Kobayashi et al., 2008; Oliver et al., 2006), and ColXVIII to assay putative changes in the NPCs. At E11.5 only few Wnt4+ cells were depicted in the WT and the ColXVIII-deficient kidney rudiments (Fig. 6 – figure supplement 1), but at E13.5, in the Six2+ cell CM population the *Wnt4^Cre^* derived floxed YFP signal was clearly detected in close proximity of the UB tip region in the NPCs in both genotypes (Fig. 6).

**Figure 6.**
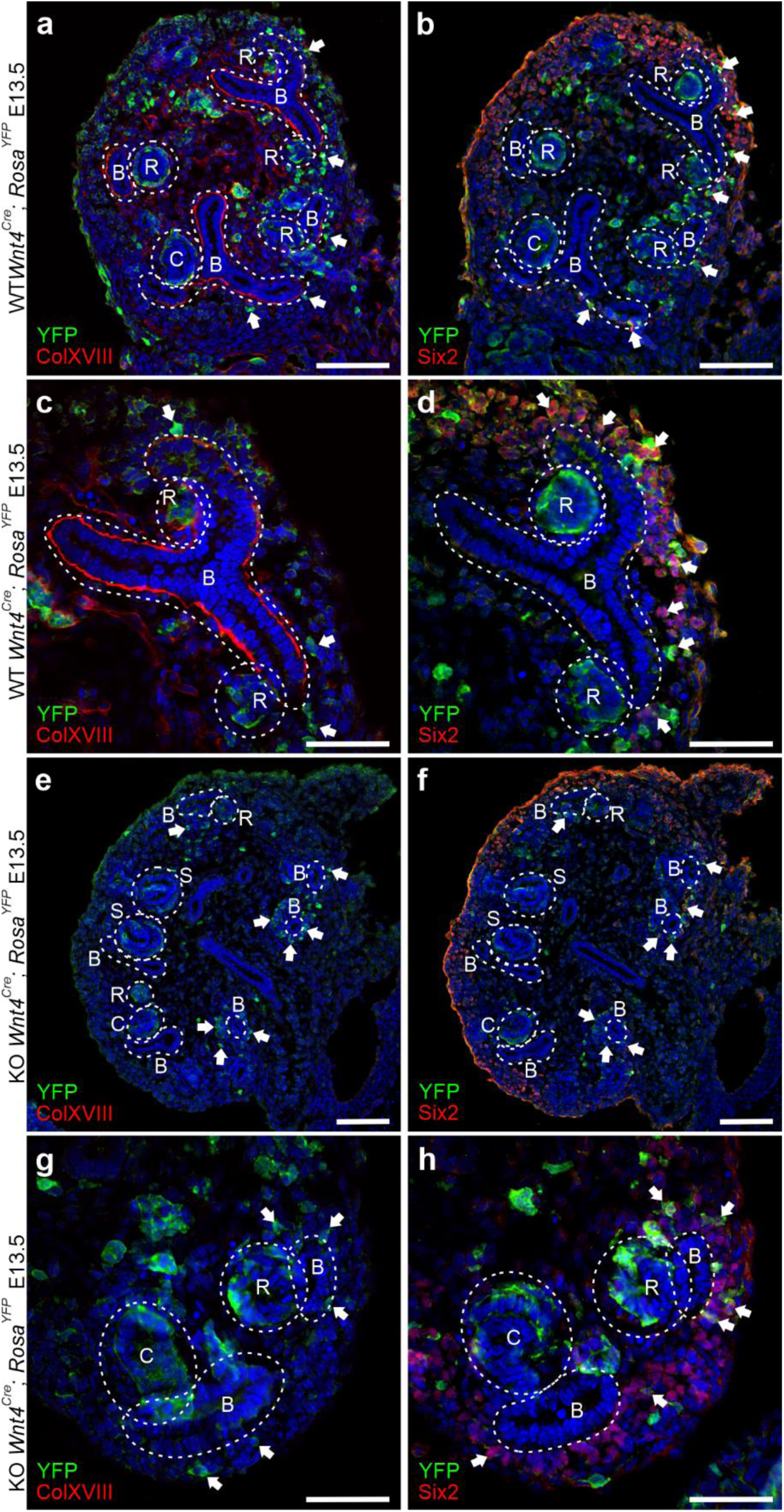
Wnt4-positive cells were detected in NPC population and in a close proximity of the ColXVIII expression in E13.5 kidneys. (a) In the WT E13.5 kidneys, the Wnt4+ cells (YFP-positive, arrows) were detected around the branching tubules (B) in the NPC population and in the renal precursors (renal vesicles (R) and comma- (C) and S-shape (S) bodies) closely localised with ColXVIII expression which was detected in the BMs of the branching tubules and the renal precursors. (b) Some of the Wnt4+ cells were also positive for Six2 indicating that these cells present NPCs. Some of the Six2 and Wnt4 positive cells are indicated with arrows. (c) A magnification of one branch surrounded by Wnt4+ cells (arrows) and Wnt4+ renal vesicles. (d) A magnification of the same branch indicating that some of the Wnt4+ cells express also Six2 and represent NPCs (arrows) in the cap mesenchyme (CM) area. (e) E13.5 ColXVIII KO kidneys stained against YFP and ColXVIII showed that the Wnt4+ cells (arrows) localised around the branching tubules and renal precursors similarly with the WT kidneys. (f) In the KO kidneys, some of the Wnt4+ cells expressed also Six2 (arrows) in the CM as seen in the WTs. (g) A magnification of a CM area of the E13.5 KO kidneys, where Wnt4+ cells were detected in the renal vesicles (R), comma-shape body (C) and in the CM area around the branches (B). (h) A magnification of the same area than in (g) stained against Six2 and YFP indicated that some of the Wnt4+ cells express also Six2 (arrows) in the CM area. Bar(a, b, e, and f): 100 μm and bar(c, d, g, and h): 50 μm.

Since the data suggested that ColXVIII may impact the NPCs and that its expression was found in the ureteric tip BMs lining the NPC population, we went on to later developmental stages and stained the E15.5 kidneys as whole-mount with the NPC biomarker Six2 and the tubular epithelial marker TROMA-1 (Keratin-8) (Djudjaj et al., 2016; Skinnider et al., 2005). From multiphoton microscope imaged kidneys, each ureteric tip-cap domain, indicating individual nephrogenic niches, were separately analysed (Fig. 5f, Movie 3). This revealed that in the KO embryonic kidneys and those that lack specifically the short (P1) ColXVIII isoform, or alternatively the two longer ColXVIII isoforms (P2) all had less Six2+ cells in foci of nephrogenic niches when compared with WT embryos (27%, 36% and 34% less, respectively, Fig. 5g).

Since we found fewer Six2+ NPCs in all ColXVIII mutant mice at E15.5, we studied also the earlier developmental stage (E13.5) aiming to define the initial signs of deficiencies in null ColXVIII embryonic kidneys (Fig. 5h). Indeed, the NPC amount as judged by Six2+ NPCs had decreased by 39% in the E13.5 KO kidneys when compared with the WTs (Fig. 5i). The results depicted that the Six2+ NPCs are decreased in the ColXVIII null embryonic kidneys at E13.5 onwards, as was the case for the isoform specific ColXVIII mutants at E15.5.

### The lack of ColXVIII leads to branching defects, reduced nephron formation, and kidney hypoplasia

We speculated that the diminution of the NPCs may be reflected to the capacity of the UB to branch (Cebrian, Asai, D’Agati, & Costantini, 2014; Oliver et al., 2006; Xu, J., Liu, Park, Lan, & Jiang, 2014). In addition, the localisation of the ColXVIII in the BMs of the developing ureteric tree, and especially the short isoform in the BMs around the ureteric tip cells, progenitors of the collecting ducts (Kurtzeborn, Kwon, & Kuure, 2019), suggested a potential role for ColXVIII in branching morphogenesis. Thus, the ureteric tree branching was imaged at defined developmental time points by optical projection tomography (OPT). The ureteric tree was stained with TROMA-1 and ureteric tips with the Wnt11.

The OPT studies revealed in all ColXVIII mutant mouse lines in comparison to control a significant reduction in the number of terminal branch points, which reflects the number of the ureteric tips (Fig. 7a,b – figure supplement 1a). The reduction in the ureteric tip count was most notable in the kidneys of P1 mice that were deficient of the short ColXVIII isoform. The same trend was also seen in total KO kidneys and in the P2 mice lacking the two longer isoforms (Fig. 7b). The OPT analyses of Wnt11 expression of mutant embryonic kidneys and controls at E14.5, E16.5, and newborns (NB) also demonstrated a reduction in the number of the ureteric tips from E14.5 onwards until birth in all ColXVIII mutants (Fig. 7 – figure supplement 1b-h) with some variation in the case of the fetuses lacking the two longer isoforms (Fig. 7 – figure supplement 1b-h). Specifically, the bifurcational branching events and ureteric tree length were reduced in all mutant mice when compared with controls at E15.5 (Fig. 7c,d). The kidneys of all mutant mice also had a slightly increased branching angle between the root branch and extending branch points, reflecting sparser branching of the ureteric tree (Fig. 7e).

**Figure 7.**
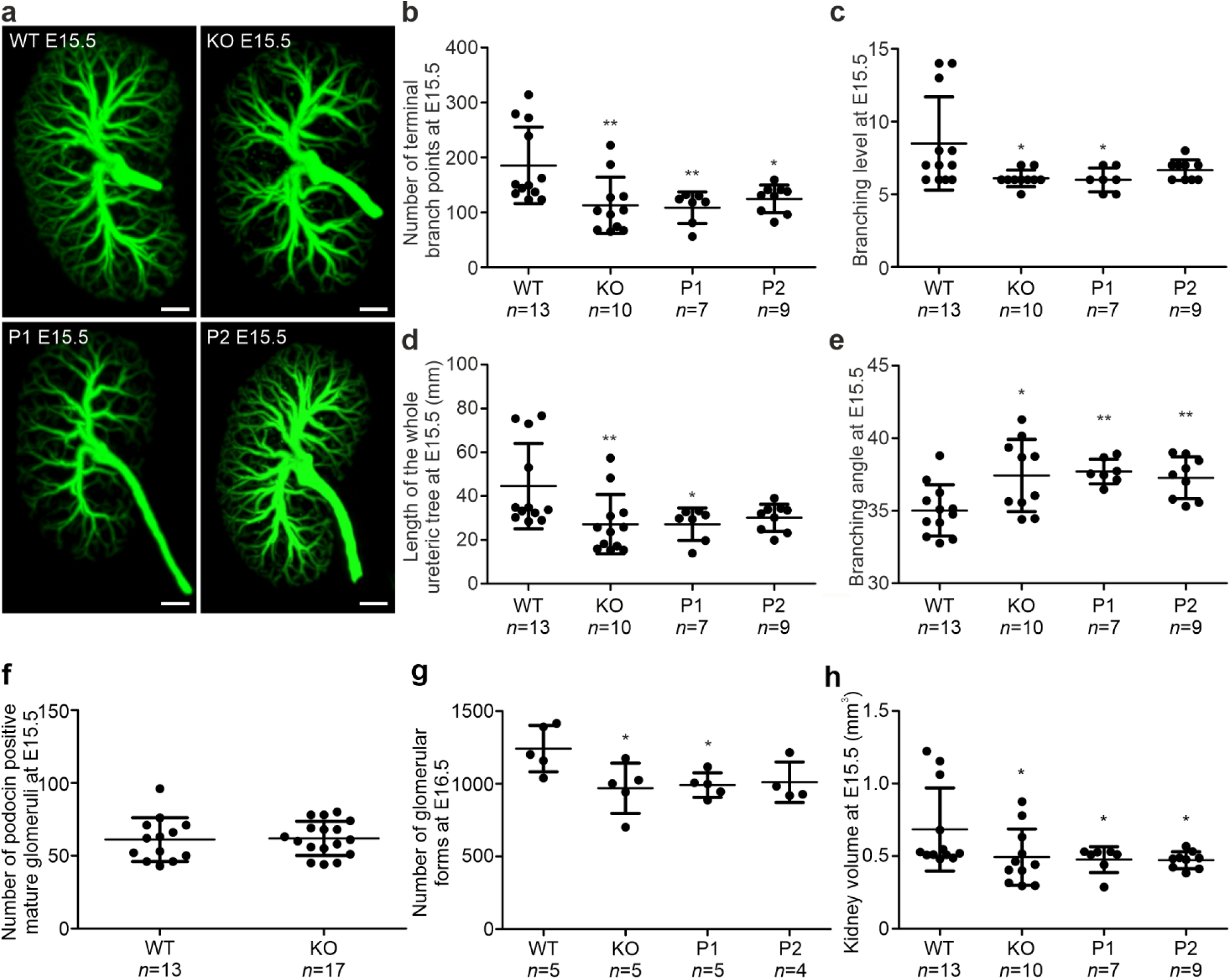
Lack of ColXVIII and its three isoforms led to branching defects during renal development. (a) The OPT images of TROMA-1 stained E15.5 kidneys suggested a reduction in the branching of the ureteric tree in kidneys lacking all ColXVIII forms (KO) or the short form (P1). N(WT)=13, n(KO)=10, n(P1)=7, n(P2)=9. Bar: 200 µm. (b) The number of terminal branch points was reduced at E15.5 in all ColXVIII mutant kidneys (KO, P1, and P2) when compared with WT kidneys. (c) Also, the branching level, which indicates the number of bifurcational branching events, was significantly reduced in kidneys lacking all or specifically the short form (KO, P1). (d) The length of the ureteric tree was significantly reduced in the E15.5 ColXVIII KO and P1 kidneys in comparison with WT kidneys. (e) The branching angle, which measures the angle between the root branch and extending branch points, was larger at E15.5 in ColXVIII mutant kidneys (KO, P1 and P2) than in WT kidneys. (f) Lack of ColXVIII did not change the number of podocin-positive mature glomeruli detected by OPT at E15.5. N(WT)=13, n(KO)=17. E15.5 was selected for podocin staining since the antibody did not penetrate into the older kidneys. (g) The number of glomeruli and glomerular precursors was decreased in all ColXVIII mutant mice (KO, P1 and P2) compared with WT mice at E16.5, but the difference was significant only between embryos lacking all or specifically the short form (KO, P1). N(WT)=5, n(KO)=5, n(P1)=5, n(P2)=4. E16.5 time point was used for calculations to take more of all the different glomerular forms into account. (h) ColXVIII mutant embryos (KO, P1, and P2) had smaller kidneys compared with WT embryos. The graph shows the E15.5 time point from TROMA-1 OPT analysed kidneys and figure supplement 1 E14.5 and E16.5 kidney volumes. In graphs, mean±s.d. is presented. *P<0.05, **P<0.01 (Mann-Whitney U -test).

To analyse when the branching defect begins in the ColXVIII-deficient mice, the E11.5, E12.5 and E13.5 WT and KO kidneys were stained against TROMA-1 and imaged either with confocal microscopy (E11.5 and E12.5) or OPT (E13.5) (Fig. 8a,c,e). The results showed that the branching of the tubular tree from E11.5 to E13.5 was comparable between the WT and KO kidneys (Fig. 8b,d,f).

**Figure 8.**
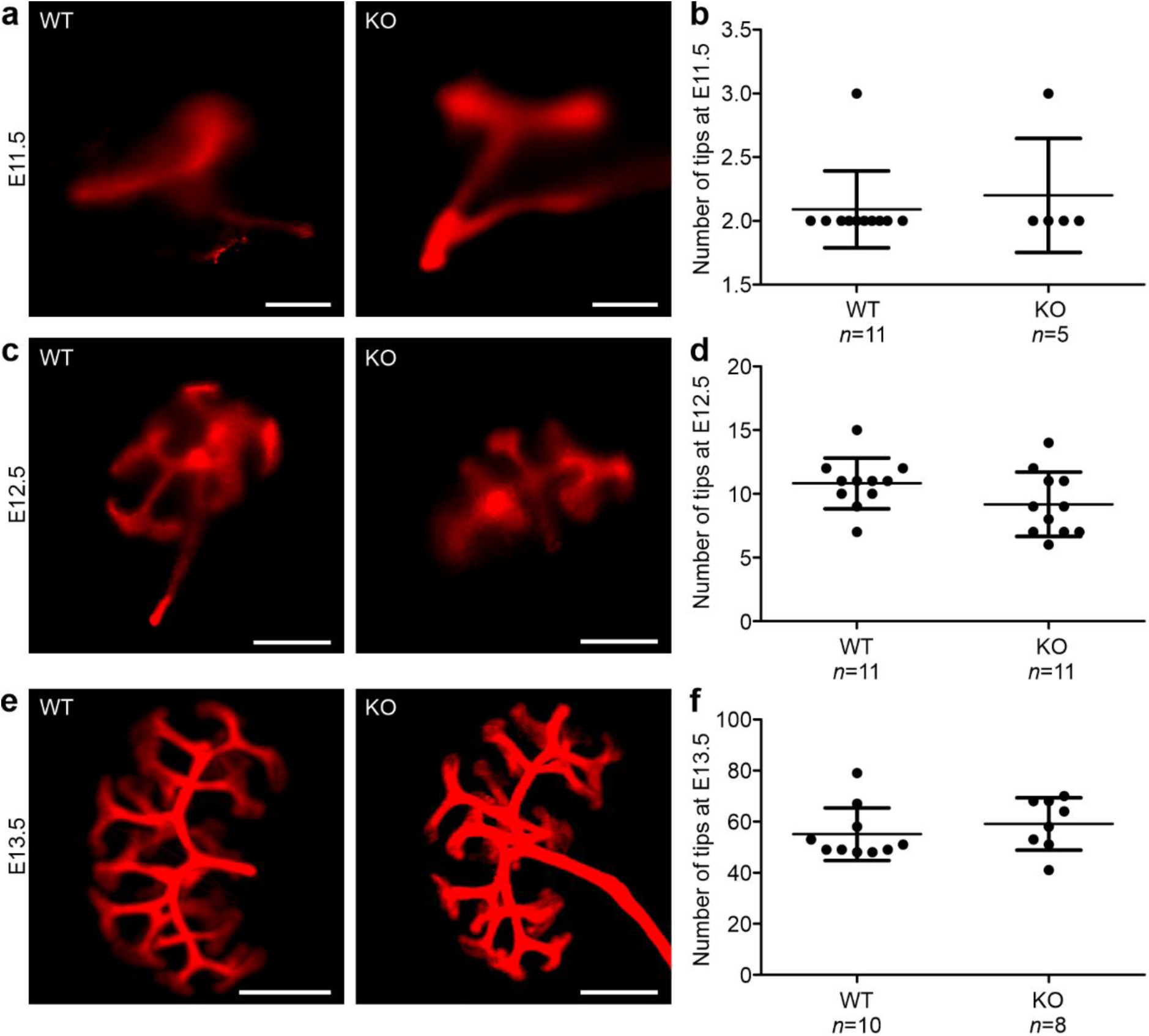
The branching defect in ColXVIII mutants begins after E13.5. Representative images of E11.5 (a), E12.5 (c) and E13.5 (e) WT and ColXVIII KO kidneys. The number of ureteric tips was comparable between the WT and the KO kidneys at E11.5 (b), E12.5 (d) and E13.5 (f). In graphs, means ±s.d. are shown. Bar(a, c): 500 μm, bar(e): 200 μm.

The tubular branching is connected to nephrogenesis (Cebrian et al., 2014; Cebrián, Borodo, Charles, & Herzlinger, 2004), and thus the number of glomeruli was analysed with OPT and podocin immunostaining depicting the mature glomeruli (Kirita et al., 2016; Roselli et al., 2002). At E15.5, no differences in the number of mature glomeruli between WT and KO kidneys (WT mean 61 (±35) and KO mean 61.94 (±18.06) glomeruli per kidney) were noted (Fig. 7f). Since only a few of the glomeruli are mature at this time point, the number of mature and developing glomeruli including renal vesicles, comma- and S-shape bodies, were next calculated from E16.5 serial sectioned whole kidneys. In this case, the number of glomeruli and glomerular precursors were decreased in all mutant mice, but the difference was statistically significant only for KO and P1 compared with WT (Fig. 7g).

Previous studies have linked the reduction of NPCs and tubular branching with renal hypoplasia (Cebrian et al., 2014; Cebrián et al., 2004; Chi et al., 2004; Xu, J. et al., 2014). Indeed, evaluation of the volumes of E14.5, E15.5, E16.5 and NB kidneys using either TROMA-1 or Wnt11 markers and OPT revealed that from E14.5 onwards the P1 kidneys were hypoplastic and from E15.5 onwards also the KO kidneys were hypoplastic when compared with the WT (Fig. 7h - figure supplement 2). The P2 kidneys lacking only the two longer isoforms were significantly smaller than the WT kidneys only at E15.5 and NB (Fig. 7h – figure supplement 2). Thus, the decrease of the renal volume seemed to follow the observed decrease in the branching in the ColXVIII mutant mice as hypothesised. These findings indicated that ColXVIII, and especially its short form, is important for the branching of the ureteric tree, nephron formation, and overall kidney growth.

### The lack of ColXVIII leads to changes in the cell cycle progression of the Six-positive NPCs

Two differently behaving Six2+ cell populations exist, those having a fast cell cycle and are committed to differentiate, and those having a slower cell cycle and maintain the NPC population (Short et al., 2014). The change in the balance of these two Six2+ subpopulations might explain the observed decrease in the Six2+ cell number in the ColXVIII mutant kidneys (Fig. 5g,i). Thus, we used flow cytometry to analyse whether the cell cycle progression of the Six2+ NPCs would differ between WT and mutant mice at E13.5, E15.5 and E17.5 time points. At E13.5, the Six2+ cells of the KO kidneys were significantly less often in the G1 phase and significantly more often in the G2-M phase when compared with the WTs (Fig. 9a,c – figure supplement. 1a,b). In contrast, at E15.5 and E17.5, there were significantly more Six2+ cells in the G1 phase (Fig. 9d,g, - figure supplement 1c-f) and significantly less in the G2-M phase (Fig. 9f,I – figure supplement 1c-f) in the KO kidneys compared with WT kidneys. No significant changes were detected in the S-phase (Fig. 9b,e,h)

**Figure 9.**
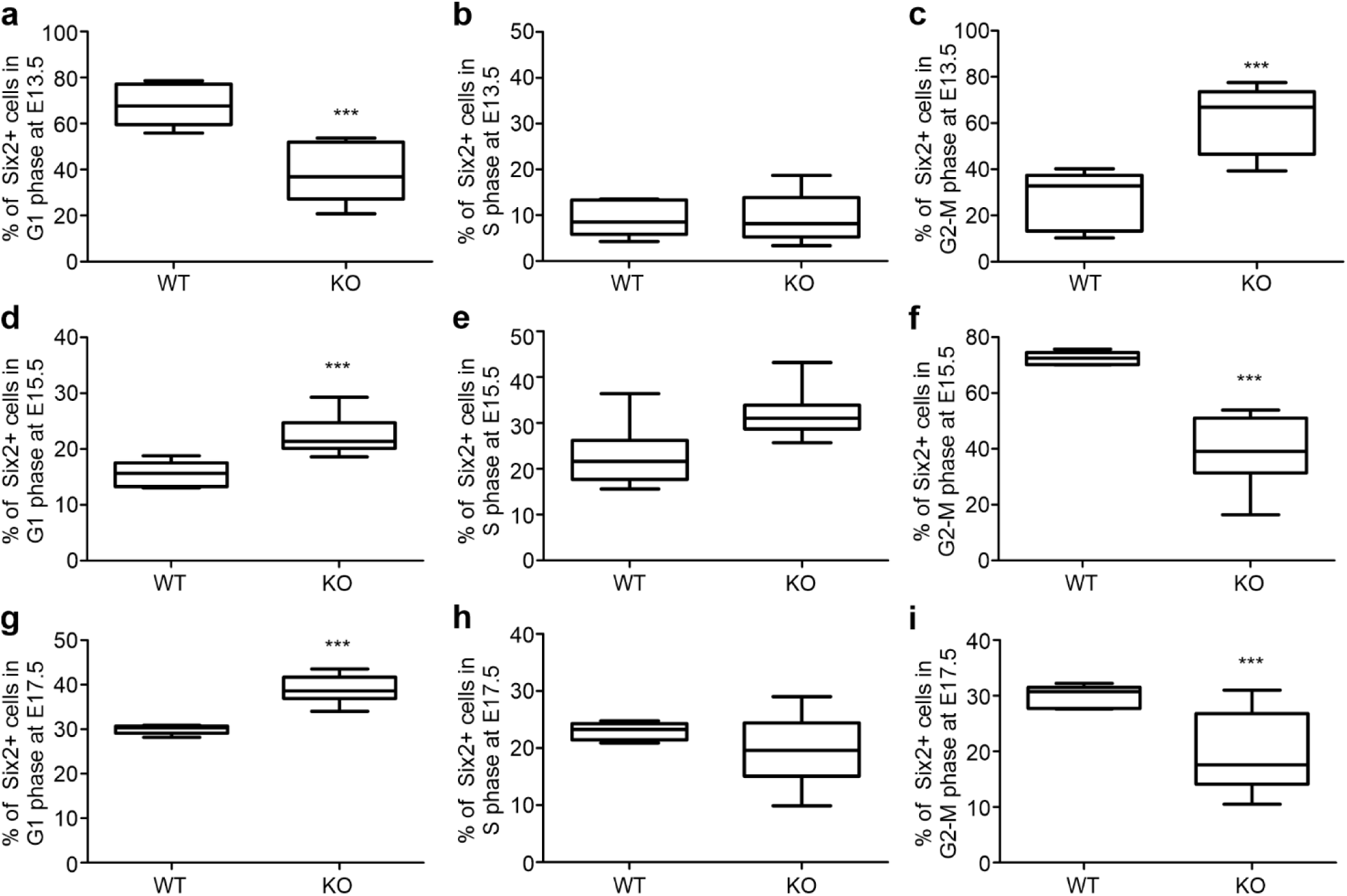
ColXVIII deficiency changed cell cycle progression during kidney development. (a) At E13.5, the number of Six2+ cells were significantly less in the G1 phase in the KO kidneys compared with the WTs. No change was detected in the S phase (b) between the WT and KO kidneys, but the Six2 cells were significantly more in G2-M phase in the KOs at E13.5 (c). At E15.5 the number of Six2+ cells were significantly more in the G1 phase (d) and significantly less in the G2-M phase (f) in the KO kidneys when compared with the WT but the difference in the S phase (e) was not significant. Similarly, at E17.5, the number of Six2+ cells was significantly higher in the KO kidneys in the G1 phase (g) and significantly lower in the G2-M phase (i) in comparison with the WT kidneys. No difference was detected in the S phase (h). N(WT E13.5)=4, n(KO E13.5)=7 litters analysed separately. N(WT E15.5)=6, n(KO E15.5)=11, n(WT E17.5)=5, n(KO E17.5)=6 independent acquisitions in flow cytometry analysis per sample collected from three different litters. In graphs, the boxes show 1st and 3rd quartiles with mean, and the whiskers indicate min and max values. ***P<0.0001 (Mann-Whitney U -test).

We next analysed the number and localisation of the Ki67-positive (Ki67+) proliferating cells within the Six2-positive NPC cells using immunostaining against Ki67, Six2 and tubular marker TROMA-1. At E13.5, the KO kidneys had less Ki67+ cells in the Six2+ NPC population when compared with the WT (Fig. 10a,c). The proliferating Ki67+ cells localised less often to the cortex side of the nephrogenic niche (Fig. 10a, white arrows, d) where reside the slow cycling NPCs which maintain the NPC population. Similarly with the E13.5 timepoint, there were significantly less Ki67+ cells in the Six2+ population at E16.5 in the KO kidneys when compared with the WTs (Fig. 10b,e), but no difference was observed in the localisation of the Ki67+ cells in the niches (Fig. 10f). Thus, the lack of ColXVIII probably affects the ratio of the fast versus slow cycling Six2+ cells, where at E13.5 the KO kidneys have more fast cycling Six2+ cells locating at the medullar side of the nephrogenic niche that are committed to differentiate. This fastened differentiation, in turn, might be a cause for the overall reduction of the Six2+ cells in niches observed at E13.5 onwards. At E15.5 and E17.5 the remaining Six2+ population in the KO kidneys represent more the slow cycling NPCs, maintaining the NPC population which is crucial for the continuity of the renal development.

**Figure 10.**
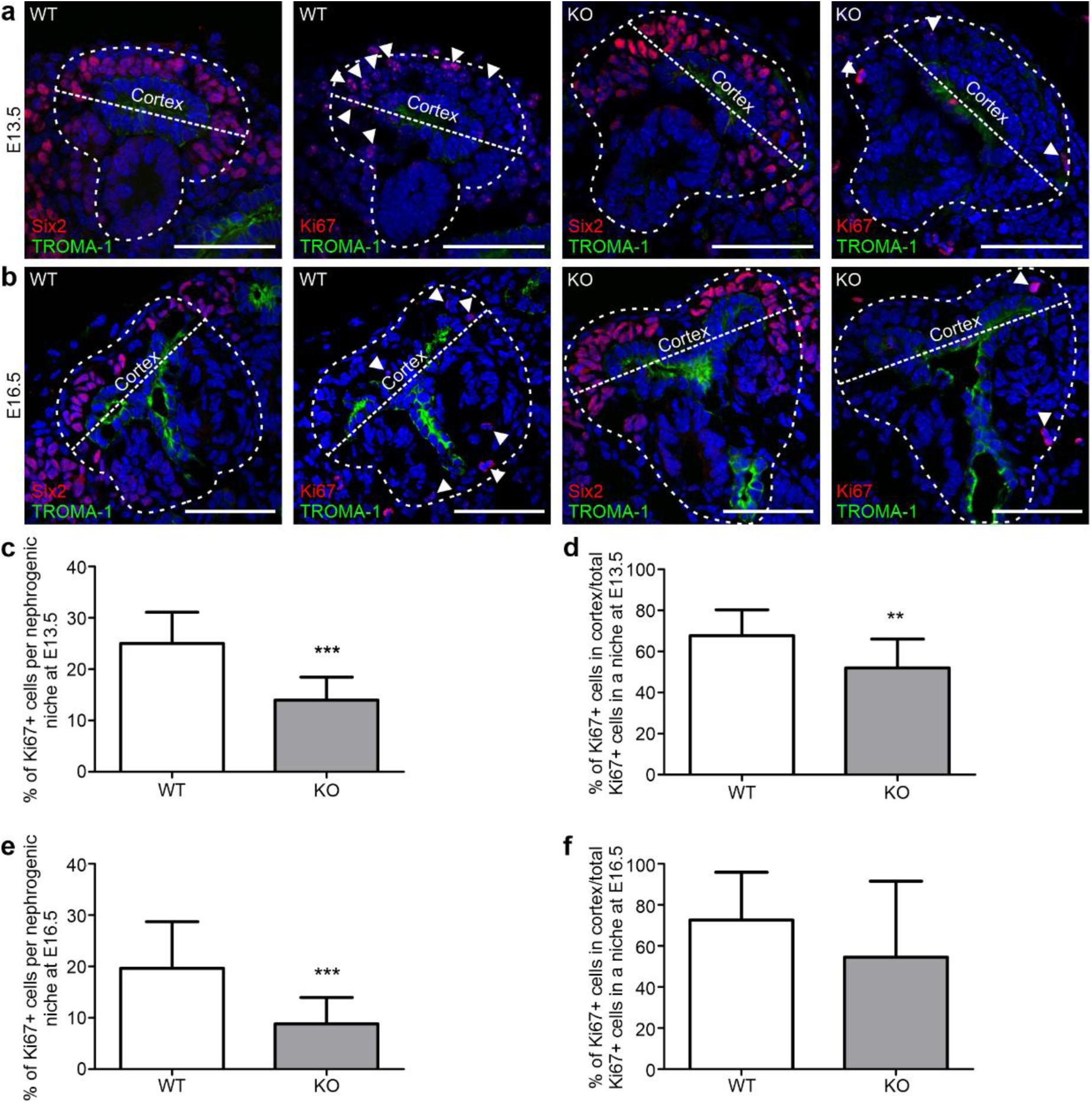
The lack of ColXVIII changes the behaviour of the proliferating Six2+ cells. (a) Images from the E13.5 kidneys stained against Six2, Ki67 and TROMA-1 indicated that the WT kidneys had more proliferating Six2+ and Ki67 positive (Ki67+) cells than the KO kidneys, and that these Ki67+, Six2+ cells localised mainly to the cortex side of the niche. (b) Similarly than at E13.5, at E16.5 the WT kidneys seemed to have more of the proliferating Ki67+, Six2+ cells in the nephrogenic niches when compared with the KO kidneys. Arrowheads point to Ki67+ cells in the nephrogenic niche indicated by the dashed lines. Bars: 50 μm. (c) The calculations of the Ki67+ cells from each niche showed that the KO kidneys had indeed significantly less Ki67+ cells at E13.5. The graph shows the ratio of the Ki67+ cells/total number of Six2+ cells in each counted niche. (d) In the KO kidneys the Ki67+, Six2+ cells also localised significantly less to the cortex side of the niche. (e) At E16.5 the KO kidneys had less proliferating cells in the niches when compared with the WT. (f) However, at E16.5 the localisation of the Ki67+ cells in the niches was comparable between the WT and KO kidneys. In graphs, columns indicate mean±s.d. N=3 kidneys in each genotype and timepoint. **P<0.01, ***P<0.0001 (Mann-Whitney U -test).

### The N-terminus of ColXVIII rescues the branching defect in ColXVIII-deficient kidneys

The N-terminal domains of ColXVIII have been reported to have functions in other tissues, which suggests that they might also participate in the regulation of tubulogenesis. For example, the Frizzled domain of ColXVIII has been shown to inhibit Wnt/β-catenin signalling, and the glycoprotein TSP1, resembling the TSP1-like domain of ColXVIII, can bind to integrins (DeFreitas et al., 1995; Hendaoui et al., 2012; Quélard et al., 2008). Thus, we tested whether a recombinant N-terminal noncollagenous fragment of ColXVIII, containing the Frizzled, the TSP1-like and the MUCL-C18 domains, would exert any effect on tubular branching in kidney organ cultures (Fig. 1). Without the fragment, the branching of the ColXVIII KO kidneys stopped after a few branches, but the WT kidneys grew and branched robustly (Fig. 11a,b). In contrast, when the kidney rudiments were cultured with 500 ng/ml or 1000 ng/ml of the N-terminal fragment, the phenotype of the KO kidneys was rescued, and the branching occurred normally (Fig. 11a,b). Interestingly, the WT kidneys also seemed to benefit from the fragment by forming more branches, but the effect was moderate compared with the KO kidneys (Fig. 11b). This indicated that the N-terminal domains of ColXVIII affect tubulogenesis and strikingly rescue the branching defect of the ColXVIII-deficient kidneys.

**Figure 11.**
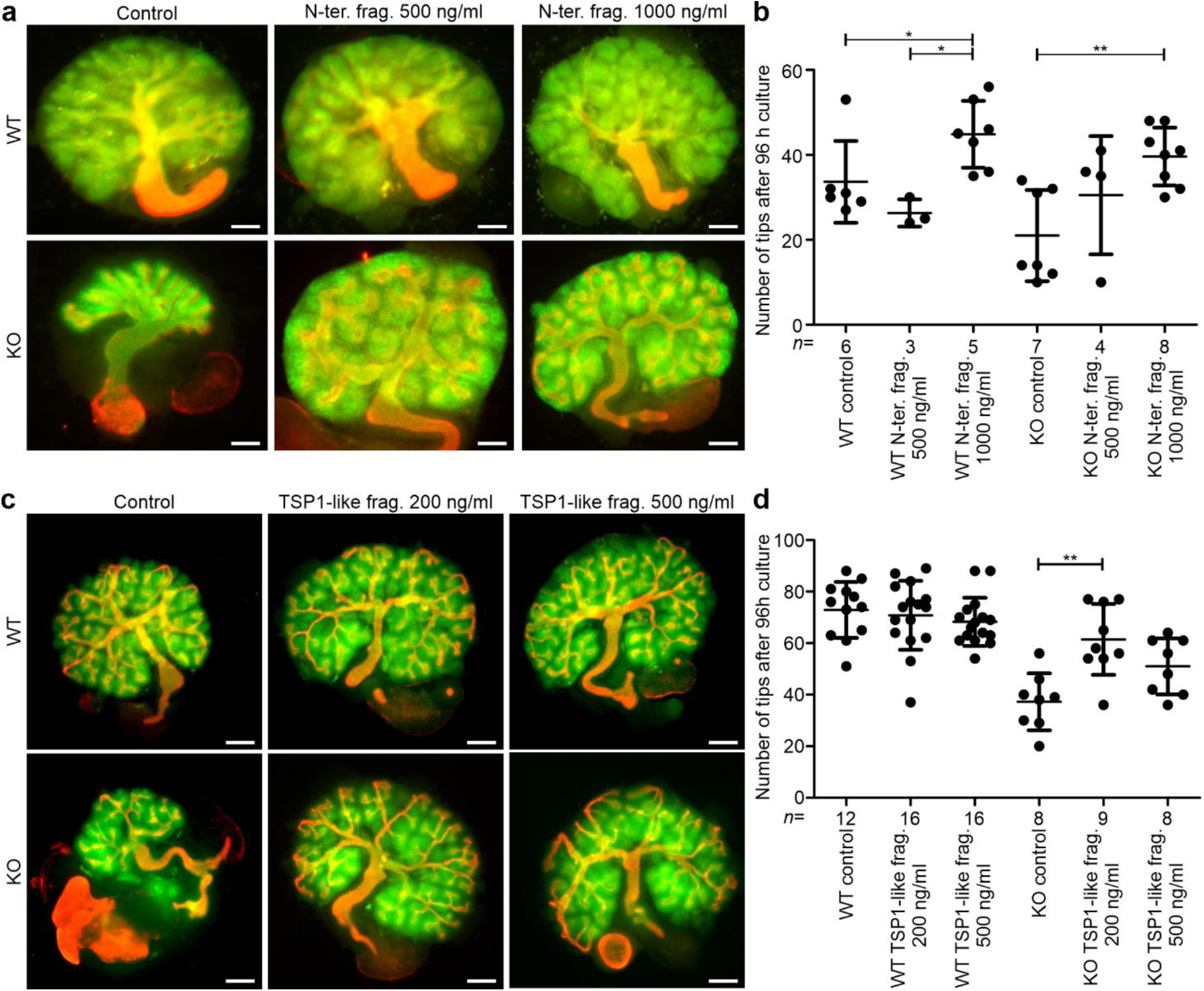
The N-terminal noncollagenous fragment of ColXVIII, and the TSP1-like domain within it, rescued the branching defect observed in the ColXVIII-deficient kidneys. (a) E11.5 WT and ColXVIII KO kidneys were cultured for 96 h in standard culture conditions or with addition of 500 ng/ml or 1000 ng/ml of the N-terminal fragment (N-ter. frag.) (see also Fig. 1). Bar: 100 µm. (b) The number of the ureteric tips of the kidneys cultured with or without the N-terminal fragment for 96 h was assessed by calculating the tips of the TROMA-1 stained ureteric tree. (c) E11.5 WT and KO kidneys cultured for 96 h in standard culture conditions or with addition of 200 ng/ml or 500 ng/ml of the TSP1-like recombinant fragment (TSP1-like frag.) (see also Fig. 1). Bar: 100 µm. (d) The number of ureteric tips after 96 h of culture with or without the TSP1-like fragment. In (b) and (d), the number (n) of cultured kidneys/condition is indicated under the column. Green: Pax2 staining indicating the mesenchyme; red: TROMA-1 staining indicating the ureteric tree. In graphs, mean±s.d. is shown. *P<0.05, **P<0.01, and ***P<0.001 (Kruskall-Wallis test with Dunn’s multiple comparison test).

The TSP1-like domain is the only common N-terminal domain for all ColXVIII isoforms. Since the branching analyses indicated that especially the short ColXVIII isoform is important for ureteric branching, we performed the kidney organ cultures with the recombinant TSP1-like fragment (Fig. 1) to evaluate its role in the branching morphogenesis. The kidney cultures were treated either with 200 ng/ml or with 500 ng/ml of the TSP1-like fragment.

The treatment of the WT kidneys with the TSP1-like fragment did not cause any changes in the branching morphogenesis when compared with the untreated kidneys (Fig. 11c,d). In contrast, the treatment of the KO kidneys with 200 ng/ml of the TSP1-like fragment caused a similar rescue effect of the ureteric branching than was seen with the full N-terminal fragment (Fig. 11c,d). The treatment with 500 ng/ml of the TSP1-like fragment also benefitted the branching of the KO kidneys but less than the treatment with less of the TSP1-like fragment (Fig. 11c,d). These results suggest a novel role for the TSP1-like domain of ColXVIII as the main branching promoting part of the N-terminus.

### Integrin α3β1 is a potential receptor to mediate the branching promoting effect of the TSP1-like domain

The TSP1 protein has been shown to bind integrin α3β1 and, interestingly, the kidney phenotypes of *Itga3*-null mice, as well as *Itgb1* knockout in UB, resemble the one described here for the ColXVIII-deficient mice (DeFreitas et al., 1995; Kreidberg et al., 1996; Wu et al., 2009; Zhang et al., 2009). Hence, the colocalisation of integrin α3 and β1 subunits, ColXVIII and Six2 was studied by immunostaining E16.5 kidney sections. The expression of both integrin subunits overlapped with the expression of ColXVIII and the Six2+ cells were detected in close proximity (Fig. 12 and 13). Moreover, lack of ColXVIII did not affect the expression patterns of the α3 and β1 integrin subunits.

**Figure 12.**
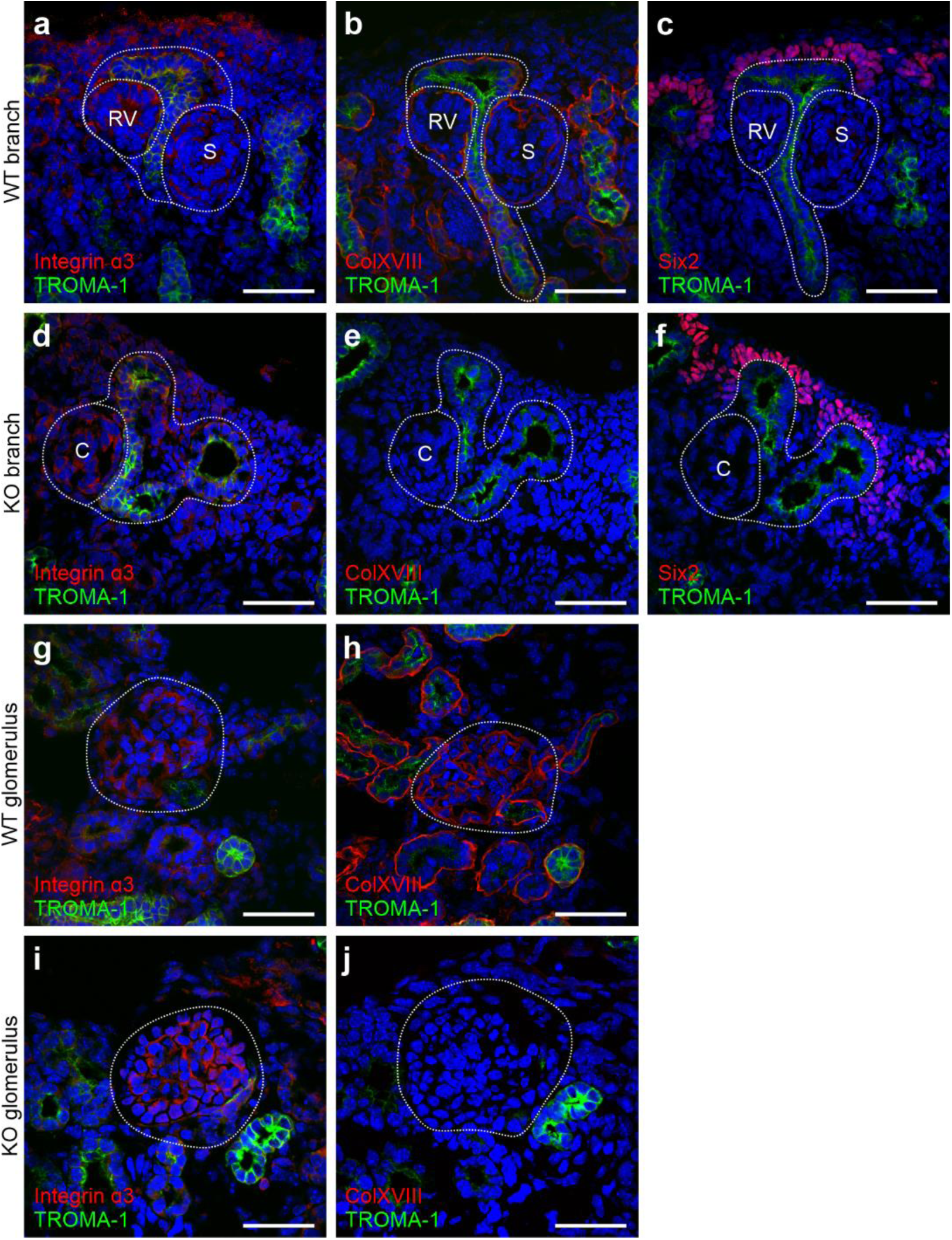
Integrin. α**3 subunit expression in E16.5 kidneys was detected in the same locations as ColXVIII and in a close proximity of the Six2-positive NPCs.** Integrin α3 staining of (a) WT and (d) ColXVIII KO kidneys in the branching tubules marked by TROMA-1 staining, and renal precursors under the branches. Dashed areas indicate integrin α3-positive branch and the renal precursors (RV=renal vesicle, C=comma-shape body, S=S-shape body) of the branches. The same branch than in (a) and (d) in the next section stained against ColXVIII and TROMA-1 in the WT (b) and the KO (e) as well as against Six2 and TROMA-1 in the WT (c) and the KO (f). Integrin α3 expression in the glomerulus (dashed area) of the WT (g) and KO (i) kidneys. ColXVIII expression was detected in same locations than the integrin α3 in the WT glomerulus (h, dashed area). TROMA-1 staining was used in the identification of the same structures in the serial sections. In the KO kidneys ColXVIII expression was not detected in the glomerulus (j). Bar: 50 μm. Green= TROMA-1, red= integrin α3/ColXVIII/Six2.

**Figure 13.**
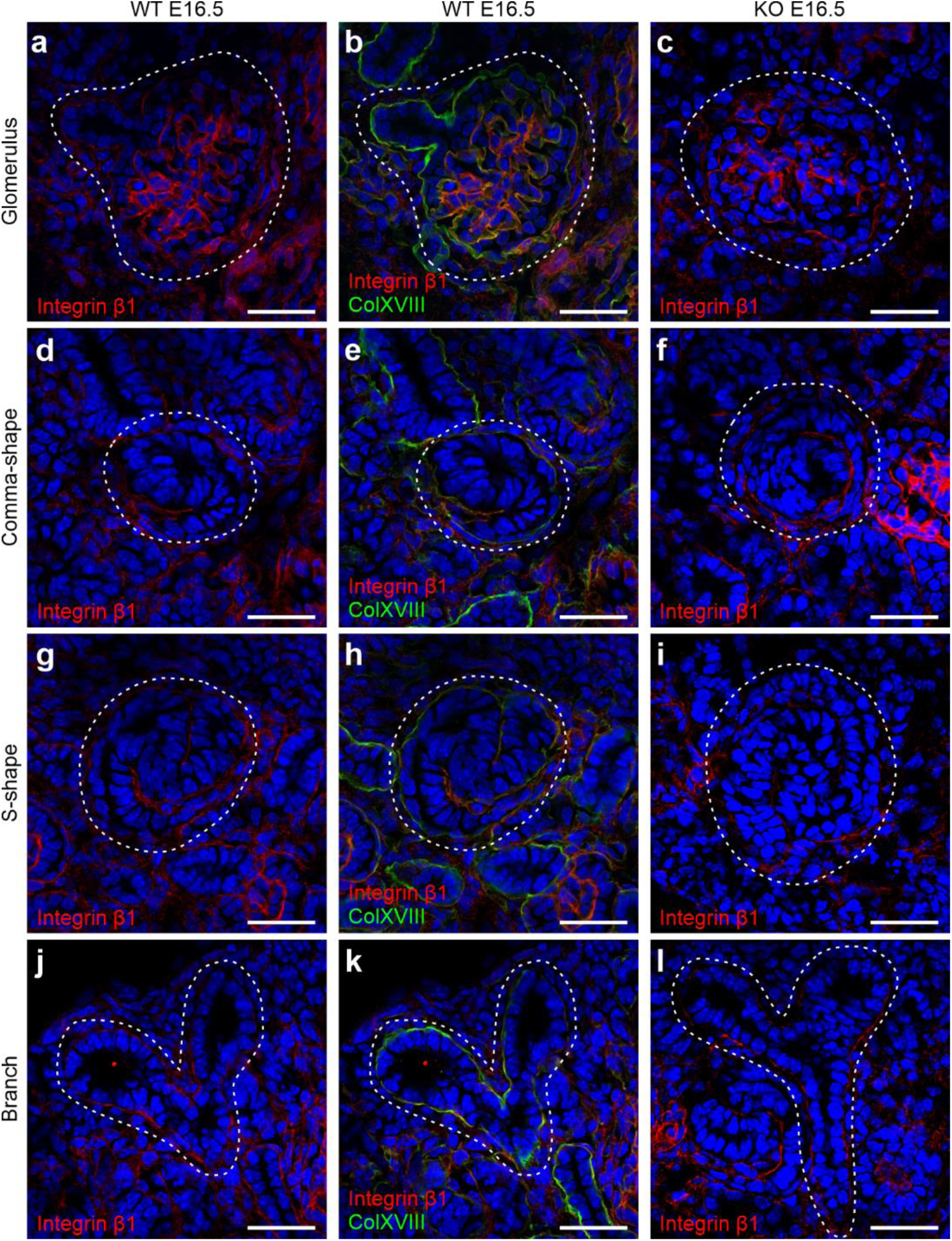
ColXVIII colocalised with integrin. β**1 in the BMs during renal development.** Images show integrin β1 (red) and ColXVIII (green) expression at E16.5, where yellowish and orange colors indicate colocalisation in the merged images. Integrin β1 expression in the glomeruli of (a) WT and (c) KO kidneys. (b) Merged image of ColXVIII and integrin β1 staining in the WT glomerular BM. Integrin β1 expression in a comma-shaped body in (d) WT and (f) KO kidneys. (e) Merged image of ColXVIII and integrin β1 staining in the WT comma-shaped body’s BM. Expression of integrin β1 in an S-shape body in (g) WT and (i) KO kidneys. (h) Merged image of ColXVIII and integrin β1 in the S-shape body in WT kidneys. Integrin β1 expression in branching tubules in (j) WT and (l) KO kidneys. (k) Merged image of ColXVIII and integrin β1 in the BMs of the branching tubules. Dashed areas indicate glomeruli, comma- and S-shape bodies, as well as branching tubules, respectively. Bar: 50 μm.

To study integrin α3β1 as a putative receptor for the N-terminal part of ColXVIII the kidney cultures were treated with integrin α3 or β1 function blocking antibodies. Inhibiting integrin α3 or β1 functions with the respective antibodies strongly inhibited the tubular branching of the cultured WT kidneys, which was seen as a reduction of the number of tips when compared with IgG-treated controls (Fig. 14a-d). In the KO cultures, the branching appeared further reduced when the integrin antibodies were added (Fig. 14a-d). The rescue of the branching defect of the KO kidneys with the N-terminal fragment (Fig. 14a-d) was inhibited by the addition of the integrin α3 or β1 blocking antibodies, and neither was there a rescue of the integrin antibody-induced branching defect of the WT kidneys (Fig. 14a-d). Thus, the results suggest that integrin α3β1 could mediate the function of the N-terminal fragment on the developing ureteric tree.

**Figure 14.**
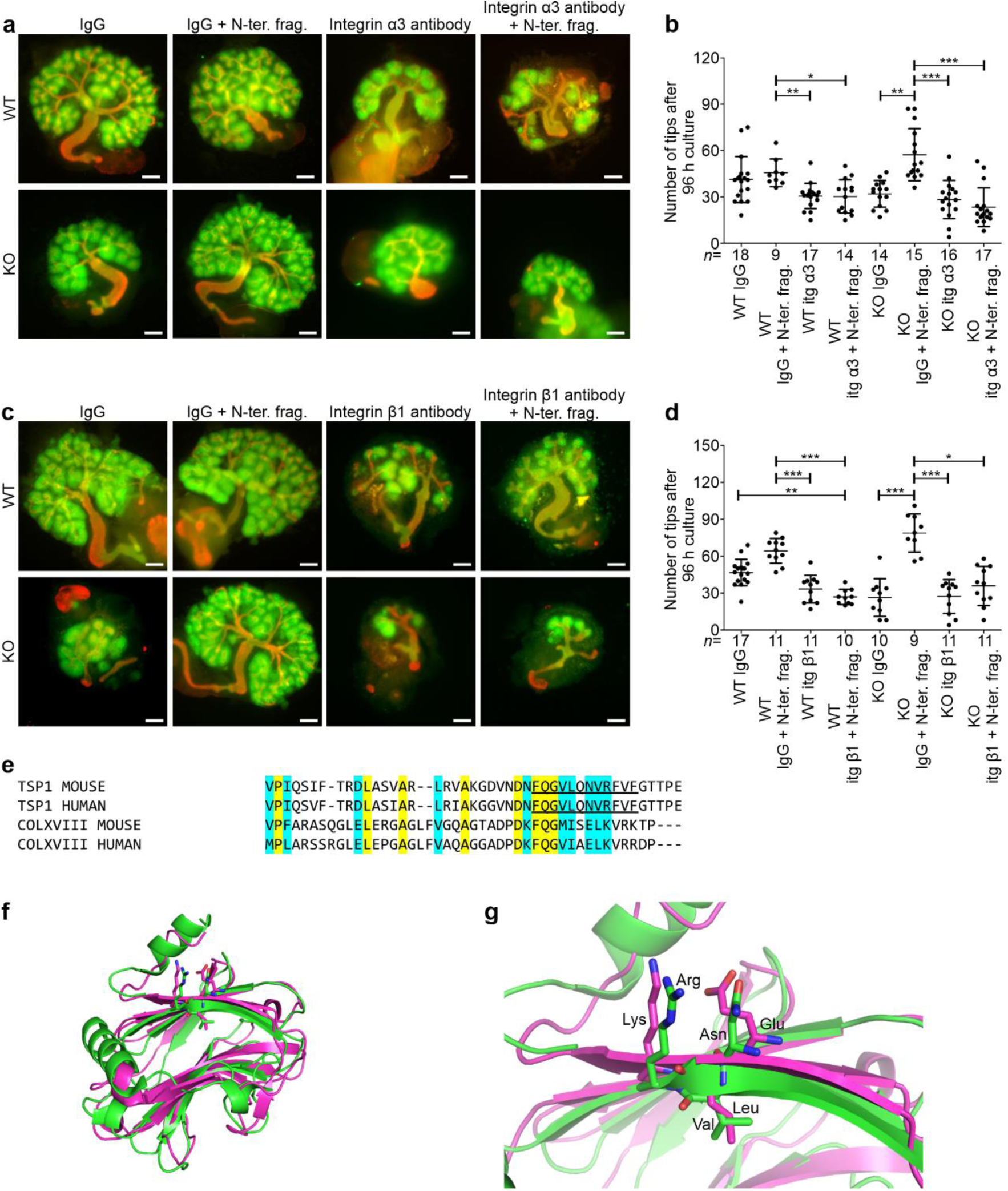
Integrin α3β1 is a potential receptor to mediate the effects of the TSP1-like domain of ColXVIII in branching morphogenesis. (a) Treatment of the WT and KO kidney cultures with an integrin α3 antibody or with a control IgG. In addition, 1000 ng/ml of N-terminal fragment (N-ter. frag.) were added to part of the cultures. Bar: 100 µm. (b) The number of the ureteric tips at 96 h of kidneys cultured with or without the integrin α3 antibody and N-terminal fragment assessed by calculating the tips of the TROMA-1 stained ureteric tree. (c) Treatment of the WT and KO kidney cultures with an integrin β1 antibody or with a control IgG. In addition, 1000 ng/ml of N-terminal fragment were added to part of the cultures. Bar: 100 µm. (d) The number of the ureter tips at 96 h of kidneys cultured with or without the integrin β1 antibody and N-terminal fragment assessed by calculating the tips of the TROMA-1 stained ureteric tree. In (b) and (d) the number (n) of cultured kidneys/condition is indicated below of the column. Green: Pax2 staining indicating the mesenchyme; red: TROMA-1 staining indicating the ureteric tree. (e) Multiple sequence alignment of the integrin α3β1 recognition sequence of mouse and human TSP1, and respective sequence of mouse and human ColXVIII. Conserved amino acids are colored in yellow and similar amino acids in cyan. The integrin α3β1 binding site FQGVLQNVRFVF of TSP1 is underlined. (f) Superimposed human TSP1 (PDB code: 1ZA4, green cartoon representation) and the homology model of mouse ColXVIII TSP1-like domain (pink cartoon representation). (g) Close view of the Asn-Val-Arg motif of TSP1 (green carbon atoms) and homologous Glu-Leu-Lys motif of ColXVIII (pink carbon atoms). In graphs, mean±s.d. is shown. *P<0.05, **P<0.01, and ***P<0.001 (Kruskall-Wallis test with Dunn’s multiple comparison test).

To study the TSP1-like domain of ColXVIII in more detail, we performed a sequence alignment and protein structure prediction. The consensus recognition site for integrin α3β1 is residues 190-201 of TSP1 (FQGVLQNVRFVF) where the sequence Asn-Val-Arg (NVR) is shown to be essential (Krutzsch, Choe, Sipes, Guo, & Roberts, 1999). Here we discovered that TSP1 and ColXVIII have conserved integrin α3β1 recognition sequences in humans and mice: out of 12 amino acid residues of the consensus sequence FQGVLQNVRFVF, ColXVIII (FQGMISELKVRK) has three identical and five similar ones as TSP1 (Fig. 14e). Although ColXVIII does not possess the essential sequence Asn-Val-Arg, the respective sequence Glu-Leu-Lys in ColXVIII is structurally comparable.

To explore the 3D structure of the binding site further, we generated a homology model of the TSP1-like domain of ColXVIII with PHYRE2 (Kelley, Mezulis, Yates, Wass, & Sternberg, 2015). Superimposing the homology model with the human TSP1 crystal structure (PDB code 1ZA4) (Tan et al., 2006) revealed a high structural similarity in the integrin binding region, thus proposing integrin α3β1 as a potential a receptor for the TSP1-like domain of ColXVIII (Fig. 14f,g).

## DISCUSSION

In this study, we found that ColXVIII is crucial in the regulation of the NPC behaviour and that ColXVIII deficiency leads to altered tubular branching, reduced nephron formation, and kidney hypoplasia. In addition, our results indicated different expression patterns of ColXVIII isoforms in the developing kidney and identified a novel role for the ColXVIII N-terminal TSP1-like domain as a driver of tubular branching which probably functions through integrins (Fig. 15).

**Figure 15.**
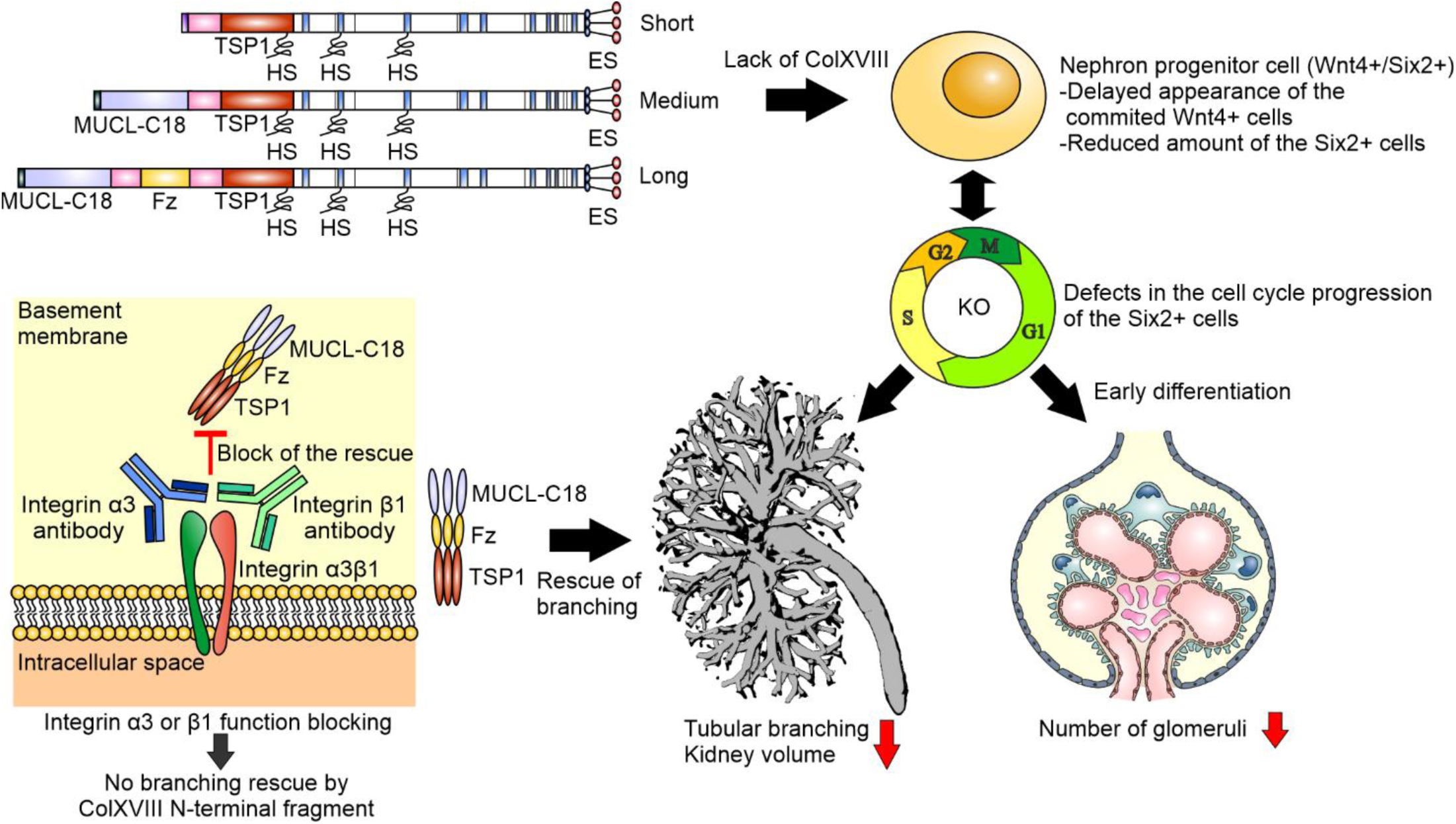
A schematic picture of the role of ColXVIII in kidney development. Lack of ColXVIII, and especially its short form, caused changes in NPC behaviour. These changes included the delayed appearance of Wnt4+ signal and changes in the cell cycle progression of the Six2+ NPCs. Furthermore, the changes in the NPC behaviour caused a defect in the branching morphogenesis of the tubular tree. This branching defect of ColXVIII-deficient kidneys can be rescued by the addition of N-terminal domain of ColXVIII *in vitro*. In addition, the results indicated that the ColXVIII N-terminal TSP1-like domain promotes the branching, and that the blocking of the function of integrin α3 or β1 could inhibit the rescue effect of the ColXVIII N-terminal fragment. ES: endostatin; HS: heparan sulphate; TSP1: thrombospondin 1-like domain; MUCL-C18: mucin-like domain in ColXVIII; Fz: Frizzled domain.

The NPC population is crucial for kidney formation as it produces inductive signals for the UB to invade the MM and to branch, while also being the cell pool from which nephrons arise (Costantini & Kopan, 2010; Little & McMahon, 2012; Oliver et al., 2006). In addition, the UB provides signals for the MM to undergo mesenchymal-epithelial transition to generate renal vesicles that mature to glomeruli (Carroll, Park, Hayashi, Majumdar, & McMahon, 2005; Halt & Vainio, 2014; Little & McMahon, 2012). In previous studies, reduced numbers of NPCs were found to cause decreased branching and nephron formation, finally leading to hypoplastic kidneys (Cebrian et al., 2014; Muthukrishnan, Yang, Friesel, & Oxburgh, 2015; Oliver et al., 2006; Xu, J. et al., 2014; Xu, P. et al., 2014). For example, in *Osr1-* and *Six2*-deficient embryos, both having a diminished CM cell population, the CM prematurely differentiates to form glomeruli (Oliver et al., 2006; Xu, J. et al., 2014). The premature differentiation of the CM cells affects reciprocal signalling between MM and UB, leading to decreased branching and nephrogenesis, as well as kidney hypoplasia (Costantini & Kopan, 2010; Oliver et al., 2006). Similar observations were made here in the ColXVIII-deficient fetal kidneys. The Six2+ NPC population was decreased in the ColXVIII-deficient mice at E13.5 and E15.5, and the cell cycle progression of these cells was affected in the ColXVIII mutants. We found that at E13.5 there were more fast cycling Six2+ cells, representing cells committed to differentiation, which in turn might lead to the observed decrease of the Six2+ NPC population. At E15.5 and E17.5 the KO kidneys had more slow cycling Six2+ cells than the WTs suggesting that the reduced NPC population in the KO kidneys tried to retain this population to ensure the continuity of the development. We detected a similar number of the mature glomeruli at E15.5 between the WT and the KO that could be explained by the fastened NPC differentiation at E13.5 in the KOs. However, the overall decrease in the number of Six2+ NPCs, and later a slowing down in the cell cycle probably leads to the detected reduction in the total number of the forming nephrons. The lack of ColXVIII was shown to also delay the time when the Wnt4+ cells appeared in the developing kidney that might indicate also a wider defect in the NPC induction in the ColXVIII mutants. The results also revealed that the NPC population is decreased before the branching defect begins in the ColXVIII mutant fetuses suggesting that this reduction is at least partly a cause for the branching defect.

The short ColXVIII isoform is the sole form in the BMs of branching tips where lies the progenitor cells of the entire collecting duct system (Kurtzeborn, Cebrian, & Kuure, 2018) and in the BMs of the nephron precursors. It is also the only form expressed during early kidney development, and thus the functional protein domains characterising this isoform are presumably the most crucial for kidney formation. A treatment with the C-terminal endostatin domain has been shown to inhibit the branching in kidney and lung *in vitro* (Karihaloo et al., 2001; Lin et al., 2001). However, our results indicated that the lack of ColXVIII, containing the endostatin domain, leads to reduced branching and kidney growth *in vivo* suggesting that other parts of ColXVIII regulate branching morphogenesis. Indeed, our *in vitro* studies with cultured kidney rudiments showed that the N-terminal domain of ColXVIII, consisting of a TSP1-like domain, a MUCL-C18 domain and a Frizzled domain, rescues the branching defect in the cultured ColXVIII KO kidneys and even promotes the branching of the WT kidneys. The results also suggested that the TSP1-like domain is the most important domain in this rescue effect, but further studies are needed to ensure this.

Earlier studies have shown that the glycoprotein TSP1 affects angiogenesis, tumour growth, and kidney injury (Kazerounian, Yee, & Lawler, 2008; Kyriakides & Maclauchlan, 2009; Maimaitiyiming, Zhou, & Wang, 2016). TSP1 also forms complexes with integrin α3β1 and, thus, regulates neuronal outgrowth, as well as endothelial cell proliferation (Calzada et al., 2004; Chandrasekaran et al., 2000; DeFreitas et al., 1995). The phenotype of *Itga3*-null kidneys resembles the phenotype seen here in ColXVIII-deficient kidneys showing reduced tubular branching, kidney hypoplasia, and ultrastructural changes similar to those observed in adult ColXVIII-deficient mice (Kinnunen et al., 2011; Kreidberg et al., 1996). Similarly, selective deletion of *Itgb1* in the UB leads to severe branching defects with nephron reduction and kidney hypoplasia (Wu et al., 2009; Zhang et al., 2009). In our *in vitro* studies, the function of the N-terminal fragment was inhibited by blocking the integrin subunits α3 or β1, suggesting that the effects of the N-terminal domain in the ureteric branching are mediated *via* integrin α3β1.

Previous publications have identified a consensus recognition site for integrin α3β1 in TSP1, specifically residues 190-201 (peptide 678, FQGVLQNVRFVF) (Furrer, Luy, Basrur, Roberts, & Barchi, 2006; Krutzsch et al., 1999). For the integrin α3β1 recognition, a highly conserved Asn-Val-Arg motif in this peptide is essential. However, peptidomimetic studies have revealed that some analogous of the Asn-Val-Arg motif are functional. Compared to the putative integrin α3β1 binding motif of ColXVIII (Glu-Leu-Lys), TSP1 peptide derivates Asp-Val-Arg, Asn-Leu-Arg and Asn-Val-Lys are functionally comparable to the Asn-Val-Arg motif (Furrer et al., 2006). Since the modification of Asn to acidic Asp retains functionality, it is likely that also structurally and chemically similar Glu is active in this position. These results indicate that the FQGMISELKVRK motif of ColXVIII is a putative integrin α3β1 recognition site. However, further studies are needed to reveal whether the TSP1-like domain could directly bind with integrin α3β1 or some other integrins and thus mediate its functions during renal development.

In conclusion, our results indicated a crucial role for ColXVIII in kidney development where it regulates NPC behaviour, branching morphogenesis, and nephron formation, all processes whose defects are linked to the increased probability of renal diseases and failure.

## MATERIALS AND METHODS

### Animal models

All animal experiments were approved by the Laboratory Animal Centre of the University of Oulu and carried out in accordance with Finnish legislation, the European Convention ETS No. 123, and European Directive 2010/63/EU. The generation of transgenic ColXVIII total knockout (*Col18a1^-/-^*, KO) targeting exon 30 in the *Col18a1* gene, and the promoter-specific mouse lines (*Mus musculus*, *Col18^P1/P1^*, P1, and *Col18^P2/P2^*, P2) targeting exons 1 and 3 in *Col18a1*, respectively, have been previously described (Aikio et al., 2014; Olsen et al., 2002). For lineage tracing studies floxed *Rosa26^LacZ^*, *Rosa26^YFP^*, and *Wnt4^EGFPCre^* mouse lines (Shan, Jokela, Skovorodkin, & Vainio, 2010) were crossed with the *Col18a1^-/-^* line to generate *Rosa26^LacZ^*; *Col18a1^-/-^*, *Rosa26^YFP^*; *Col18a1^-/-^* and *Wnt4^EGFPCre^*; *Col18a1^-/-^* lines. The WT, KO, P1 and P2 mice were first in the C57Bl/JOlaHsd background and the ColXVIII *in situ* hybridisations, immunofluorescence (IF) staining and qPCRs, E15.5 TROMA-1 and podocin OPT analyses, E14.5, E16.5 and part of the NB Wnt11 *in situ* hybridisations were done with these mice. On account of a change in the mouse lines used at the Laboratory Animal Centre, the background of the mice was changed during the study to C57Bl/N6Crl, and part of the NB Wnt11 *in situ* hybridisations and all Six2 multiphoton analyses, organ cultures, NPC cultures, integrin IF staining, flow cytometry analyses and the branching analyses of the E11.5-E13.5 were done with these mice. The *Rosa26^LacZ^*, *Rosa26^YFP^*, and *Wnt4^EGFPCre^* mice were in the C57Bl/NCrl background. The vaginal plug appearance of the inbred mice was used as a criterion for mating and the right age of the embryos was confirmed with the somite counting of the E10.5-E11.5 and the limb bud staging in the older embryos.

### *In situ* hybridisation

Section and whole-mount *in situ* hybridisations were performed as described earlier (Lin et al., 2001; Pietilä et al., 2016). For a DIG-labelled antisense and sense riboprobes, a vector cloned 783–1,592 bp fragment of mouse ColXVIII complementary DNA (cDNA) (GenBank NM_001109991.1) was linearised with BamHI and Xhol. The Wnt11 probe was generated from a linearised plasmid obtained as a gift from Professor Andrew McMahon (University of Southern California, USA) (Majumdar, Vainio, Kispert, McMahon, & McMahon, 2003). The hybridised whole kidneys were photographed with Olympus SZX12 microscope and Olympus PEN E-P1 digital camera (Olympus Corporation, Tokyo, Japan), and sections with Leica DM LB2 microscope by using C PLAN 4x/0.10, C PLAN 10x/0.22 and C PLAN 20x/0.40 objectives, and Leica Application Suite V4.3 (Leica, Wetzlar, Germany).

### Collagen XVIII antibody

Rabbit anti-collagen XVIII antibody was raised as follows. The RNA isolated from the mouse endothelioma cell line eEnd.2 was used to amplify reverse transcriptase PCR to obtain complementary DNA (cDNA) encoding the TSP1-like domain of mouse collagen XVIII. The amplified cDNA was inserted into the episomal expression vector pCEP-Pu, which contained in addition the signal peptide of BM-40, to transfect human EBNA-293 cells (Kohfeldt, Maurer, Vannahme, & Timpl, 1997). The recombinant TSP1-like domain was purified from conditioned medium using dimethylaminoethanol (DEAE) cellulose followed by Superose 6 molecular sieve chromatography and then used for immunising of a rabbit. The antibody was purified using the recombinant TSP1-like domain-coupled Sepharose 4B. The antibody was compared with existing ColXVIII antibodies and was found to be highly specific.

### Immunofluorescence and β-galactosidase staining

For immunostaining, paraffin and cryosections were cut to a 5- or 7-µm thickness. Rabbit anti-ColXVIII antibody (1 μg/ml) was produced and purified as described above. Primary antibodies against TROMA-1 (DSHB, 1:500), Six-2 (11562-1-AP; Proteintech, 1:200), podocin (P0372; Sigma, 1:300), Ki67 (Ab15580; Abcam, 0.25-0.5 μg/ml), integrin subunits α3 (AB1920; Millipore, 1:1000), and β1 (553715; BD Bioscience, 1:100), and GFP (NB100-1770SS, Novus Biologicals, 1:200) were used. The secondary antibodies used were anti-rat Alexa Fluor 594 (A11007; Molecular Probes, Invitrogen), anti-rabbit Alexa Fluor 488 (A11008, Molecular Probes, Invitrogen), anti-rabbit Cy3 (111-165-144; Jackson Immunoresearch), anti-rat Cy2 (112-225-167; Jackson Immunoresearch), anti-rabbit Alexa Fluor 647 (ab150079; Abcam), and anti-rabbit Cy2 (112-225-167; Rockland). Secondary antibodies were diluted 1:300 for section staining and 1:800 for whole-mount staining. DAPI (D9542, Sigma, 2,9 μg/ml) was used to stain the nuclei. The stained IF sections were examined with a Zeiss LSM700 confocal microscope by using Plan-Apochromat 20x/0.8 and Plan-Apochromat 63x/1.4 DIC Oil objectives and a Zen 2009 (black edition, Zeiss) program (Carl Zeiss, Jena,

Germany) or with a Zeiss LSM780 confocal microscope by using Plan-Apochromat 20x/0.8 DIC and i Plan-Apochromat 63x/1.4 DIC Oil objectives, and a Zen 2012 (black edition, Zeiss) program. Excitation wavelengths for DAPI was set to 405 nm, for Alexa Fluor 488 and Cy2 to 488 nm, for Cy3 to 514 nm, for Alexa Fluor 594 to 561 nm, and for Alexa Fluor 647 to 633 nm. Collected emission wavelengths were 410-495 nm for DAPI, 490-543 nm for Alexa Fluor 488 and Cy2, 557-681 nm for Cy3, 569-639 nm for Alexa Fluor 594, and 638-735 nm for Alexa Fluor 647.

β-galactosidase staining was performed with E11.5 embryos according to Shan *et al*. 2010 (Shan et al., 2010). The E11.5 stained samples were paraffinised, cut into 7-μm thick sections and photographed with a Leica DM LB2 microscope system by using by using C PLAN 4x/0.10, C PLAN 10x/0.22 and C PLAN 20x/0.40 objectives.

### Six2+ cell counting

The E13.5 (*n*=7-9) and E15.5 (*n*=2–6) Six2-TROMA-1 stained kidneys were embedded in 1 % low melting agar (50100; Lonza Group). The lateral cortex of the kidneys was imaged with a Nikon A1R MP+ multiphoton microscope (Tokyo, Japan) by using CFI75 Apochromat 25x/1.1 W 1300 objective and the NIS-Elements C-ER 4.6. (Nikon) program. Excitation wavelengths for Six2 and TROMA-1 stainings were set to 920nm and 1040nm. Collected emission wavelengths were 495nm-511nm for Six2 and 593nm LP for TROMA-1. The voxel size was set to 0.5µm(x)*0.5µm(y)*0.5µm(z). The images were deconvolved with the Huygens Professional version 17.10 (Scientific Volume Imaging, Hilversum, the Netherlands) and further processed with ImageJ/Fiji to crop each nephrogenic niche to separate areas.

The Six2+ cells were calculated from the cropped images by a program made with MATLAB R2017a (The MathWorks, Natick, Massachusetts, USA) for this purpose. The program pre-processed images using Hessian ridge enhancement (Hodneland et al., 2009) and H-minima transformation (Meyer, 1994). Ridge enhancement increased the cell membrane intensity and reduced the cytoplasmic fluorescence, while the H-minima transformation removed small dark regions inside cells. The pre-processed images were then segmented using Watershed transformation (Soille, 1999).

### OPT and branching analysis

The TROMA-1 (*n*=7–13), and podocin (*n*=13–17) stained and Wnt11 *in situ* hybridised (*n*=6–18) samples collected at least two different litters were prepared for OPT as previously described and scanned with the OPT Scanner 3001M (Bioptonics Microscopy, UK) (Chi et al., 2011; Short, Hodson, & Smyth, 2010). Excitation wavelengths for TROMA-1 and podocin were set to 540-580 nm. Collected emission wavelengths were 590-670 nm. Wnt11 *in situ* hybridized kidneys were imaged with bright field illumination. Pixel size was set to 3.5 μm for Wnt11 *in situ* hybridized kidneys, 7 μm for E15.5 TROMA-1 and podocin stained kidneys and 3.15 µm for E13.5 TROMA-1 stained kidneys. The obtained data were reconstructed with the Skyscan NRecon v1.6.9.4. software (Bruker, Kontich, Belgium) and analysed with Imaris software (Bitplane, Zurich Switzerland) as earlier reported (Chi et al., 2011; Pietilä et al., 2016; Short et al., 2010).

The following analysis were made with Imaris software (Bitplane, Zurich, Switzerland) from OPT imaged kidneys. The kidney volume calculations were made by measuring the x-, y- and z-axis of the kidneys crossing in the middle and approximating the volume of kidney as an ellipsoid. To calculate the volume of the kidney, 4/3πabc (where a=x-axis/2, b=z-axis/2 and c=y-axis/2) formula was used. The morphogenesis of the ureteric tree was assessed with the filament-tracing tool and the number of the positive ureteric tips from Wnt11 hybridised kidneys as well as the number of podocin-positive glomeruli with spot tracing-tool. From the filament data (schematic picture in Fig. 7 – figure supplement 1a), the values of branching angle B (measures the angle between the root branch and extending branch points), terminal branch points (reflecting the number of the ureteric tips), branching level (indicating the number of bifurcational branching events) and filament dendrite length (indicating the sum of all tubules counted together) were used to analyse the morphology of the ureteric tree.

Whole-mount E11.5 (*n*=5-11) and E12.5 (*n*=11) kidneys were stained against TROMA-1. E11.5 kidneys were embedded to 1% low melting agar and imaged with Zeiss AxioScope.A1 microscope (Carl Zeiss, Jena, Germany) by using Olympus UMPLFLN 10X/0.3 W objective with excitation wavelengths 540-580 nm and emission wavelengths 590-670 nm. E12.5 kidneys were imaged with an Olympus BX51WI microscope (Olympus Corporation, Tokyo, Japan) by using U-PlanFI 4x/0.13 objective with 555 nm excitation and long pass emission collection, and CellM Software (Olympus Corporation). The number of tips were counted with ImageJ/Fiji.

### Counting of glomeruli and proliferating cells

For glomerular counting, 7-µm thick sections were prepared of paraffin-embedded whole E16.5 kidneys (*n*=4– 5 kidney pairs), haematoxylin-eosin stained, and photographed with an Aperio AT 2 Leica Scanner and an Aperio Scan Scope Console program (Leica, Wetzlar, Germany) available at the Northern Finland Biobank Borealis. The glomeruli and glomerular precursors were calculated from every fifth section of the serial sectioned kidneys using QuPath v0.1.2 (The Queen’s University of Belfast, Northern Ireland). The cells positive for Ki67 were calculated from the cell population positive for Six2 in the next section which each surrounded one tip positive for TROMA1 using ImageJ/Fiji. Three different kidneys from two different litters per genotype and timepoint were used in Ki67 analyses.

### Flow cytometry

For flow cytometry, the E13.5 dams were injected intraperitoneally with 200 μl of 5-bromo-2’-deoxyuridine (BrDU) (552598, BD Pharmingen) early in the morning and the embryonic kidneys were collected 2.5 hours later. The kidneys of one litter (*n*=5-10 embryos per litter) were pooled together to present one sample. Five WT litters and seven KO litters were separately analysed. The cells were dissociated with Collagenase Type I (CLS-1, Worthington Biochemical Corporation) and Type IV (CLS-4, Worthington Biochemical Corporation), and DNase I (M0303S, New England Biolabs) in physiological buffer at + 37°C. The enzyme activity was stopped with fetal bovine serum (FBS) followed by the fixation and staining with the APC BrDU Flow Kit – Part A (552598, BD Pharmingen) according to the manufacturer’s recommendation with some exceptions. Together with BrDU, Six2 antibody was added (1:100) and incubated 40 min at room temperature. After washing, the secondary antibody goat anti-rabbit Alexa Fluor 488 was added (1:200) and incubated another 30 min. One drop of DAPI (R37606, Invitrogen) was added to the cell suspension couple of minutes before the analysis with BD FACSAria^TM^ III sorter (BD Bioscience, New Jersey, USA).

For E15.5 and E17.5 flow cytometry, kidneys from at least three different litters (*n*=6-9 embryos per litter) were passed through a cell mesh (352340; Falcon 40 µm Cell Strainer, Corning) and fixed with absolute ethanol at –20 ᵒC. To ensure that the difference in the method how the cells were dissociated at different time points would not cause false results, some E15.5 WT and KO were also enzymatically dissociated and labelled with BrDU and Six2 (Fig. 16). The results obtained by dissociating cells enzymatically or passing through the strainer were similar and the results obtained using the cell mesh are shown. The Six2 antibody and a goat anti-rabbit Alexa Fluor 647 (ab150079; Abcam) secondary antibody was used to detect the NPCs. The cell cycle analyses of the stained cells were performed with a flow cytometry, where 5-11 independent acquisitions per sample were done (FACSCalibur; BD Biosciences, Erembodegem, Belgium).

**Figure 16.**
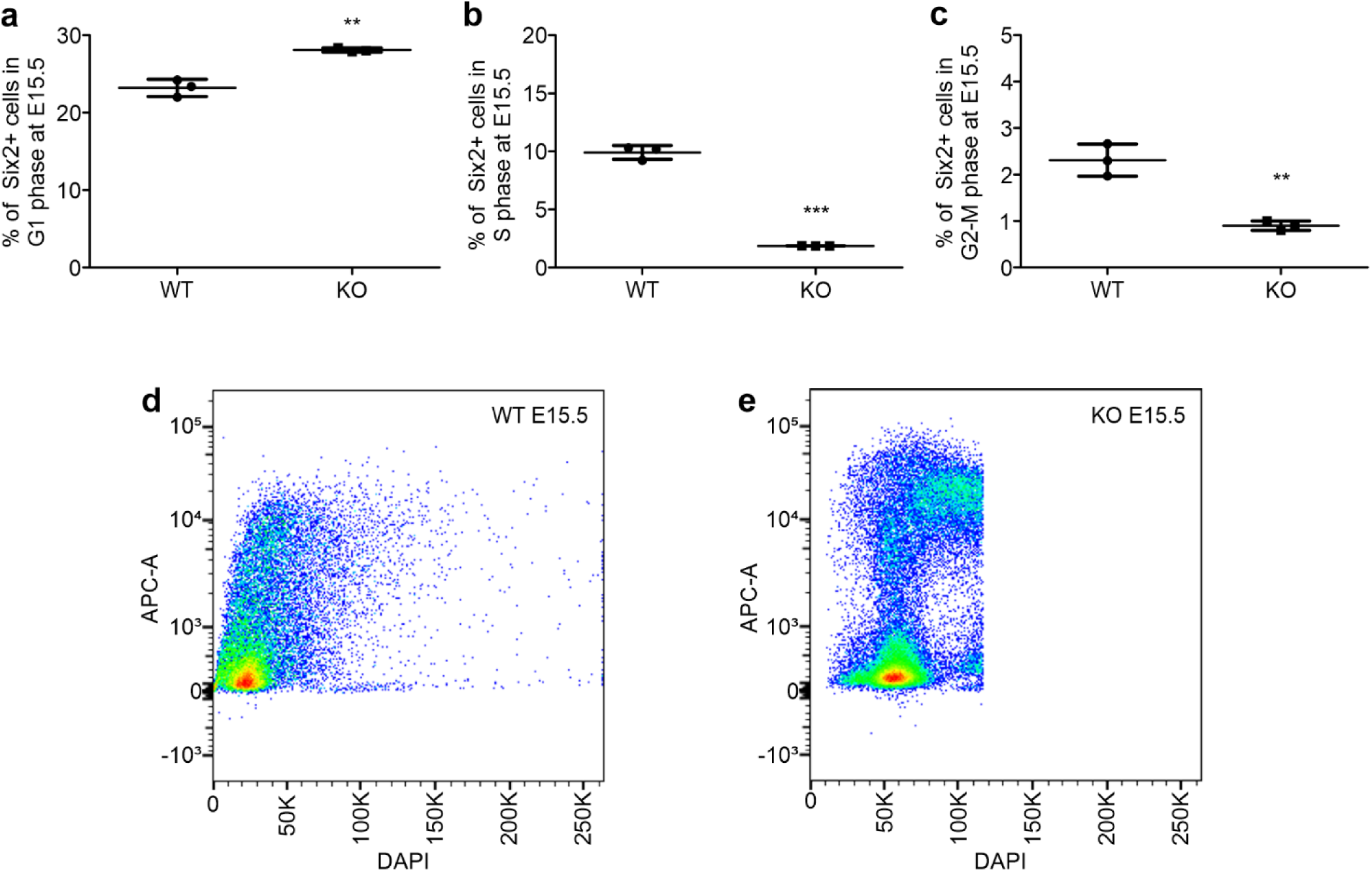
The cell dissociation method did not affect to the E15.5 cell cycle results. The enzymatic dissociation of the cells from E15.5 kidneys were labelled with BrDU and Six2. The results of the cell cycle analyses indicated that in the KOs the Six2+ cells were significantly more often in the G1 phase (a) and significantly less in the G2-M phase (c) when compared with the WTs. Similar results were detected in the analyses of the E15.5 where the kidneys were passed through the cell mesh. However, in these analyses there seemed to be significantly less Six2+ cells in the S-phase in the KOs when compared with the WTs (b). Representative images of the cell cycle analysis of the WT (d) and KO (e) kidneys. N(WT)=3, n(KO)=3 analyses, where kidneys of one litter are pooled together. In graphs, mean±s.d. is shown. **P<0,01, ***P<0,0001 (Mann-Whitney U -test).

### qPCR

RNA isolation, cDNA production and assays were performed as described (Aikio et al., 2014). The relative RNA levels were calculated with Bio-Rad CFX Manager 3.1 software using the ΔΔCq method. The used primers were for ColXVIII (recognising all isoforms) 5’-AGGACTTTCAGCCAGTGCT-3’ and 5’-AAATCTGCTCCACGGATA-3, and for ColXVIII short isoform 5’-GGATGTGCTCACCAGTTTG-3’ and 5’-CATCGATTTGTGAGATCTTC-3’. For normalisation, primers for 18S (5’-GCAATTATTCCCCATGAACG-3’ and 5’-GGCCTCACTAAACCATCCAA-3’) were used. *N*=14–22 kidney pairs were pooled for ColXVIII E11.5 analyses.

### Organ culture

The AGM region of E10.5-11.0 (39–47 somites) embryos (*n*=15–16) from *Rosa26^YFP^*; *Wnt4^EGFPCre^* modified WT, or *Col18a1^-/-^* mice were dissected, and YFP-positive embryos were selected for further processing. The cultures were set up as described by Saarela *et al*. using Cultrex (3445-005-01; Trevigen) (Saarela, U., Akram, Desgrange, Rak-Raszewska, Shan, Cereghini, Ronkainen, Heikkilä, Skovorodkin, & Vainio, 2017a). The time-lapse images were captured at 0.5 or 1 h intervals by the Zeiss LSM780 confocal microscope by using i LCI Plan-Neofluar 25x/0.8 W objective and a Zen 2012 (black edition) program or by a Leica SP8 FALCON by using HC PL APO 10x/0.40 objective and a LAS X v3.5.6 program (Leica Microsystems GmbH, Wezlar, Germany), which were also used to process and analyse the data. Environmental conditions (+37°C, 5% CO2) in time-lapse experiments were maintained by Okolab Bold Line on stage incubators (Okolab, Pozzuoli, Italy).

The N-terminal fragment of ColXVIII (Aikio et al., 2014) and TSP1-like recombinant fragment (Zaferani et al., 2014) were produced and purified as previously described. The full N-terminal fragment was applied to E11.5 kidney cultures at a concentration of 500 ng/ml (*n*=3–4) or 1000 ng/ml (*n*=6–8) at the beginning of the culturing, and controls (*n*=5–7) were cultured in medium without the fragment. Likewise, the TSP1-like fragment was applied to E11.5 at a concentration of 200 ng/ml (*n*=9-16) or 500 ng/ml (*n*=8-16), and controls (*n*=8-12) were cultured in medium without the fragment. The following antibodies were used for function blocking studies (*n*=9–18): 25 µg/ml of an anti-integrin α3 antibody, (MAB1952Z; Merck Millipore), 25 µg/ml of a mouse IgG1 monoclonal antibody (NCG01) (ab81032; Abcam), 5 µg/ml of a NA/LE hamster anti-mouse CD29 clone HM β1-1 (562219; BD Bioscience), and 5 µg/ml of a NA/LE hamster IgG2 (553961; BD Pharmingen).

To perform fragment cultures with E11.5 kidneys, a nucleopore filter with the kidney primordium was placed on top in a trowel grid, cultured for 96 h in medium (DMEM, 10% FBS, 1% streptomycin, penicillin), and incubated in a humidified atmosphere at 37°C with 5% CO2. To titrate the ColXVIII fragment concentration for the experiment, 25, 50, 100, 500, and 1000 ng/ml concentrations of the fragment were used in culture medium and changed every 48 h. A clear effect in the growth was detected at 500 ng/ml, and it was used together with 1000 ng/ml in the final experiments. The function blocking studies were performed similarly to the fragment cultures by adding either integrin α3 or β1 antibody alone or together with N-terminal fragment 1000 ng/ml, and for controls, adding IgG alone or with the N-terminal fragment 1000 ng/ml. After culture, the kidney primordia were fixed in 4% paraformaldehyde (PFA) and processed for whole-mount-immunostaining with TROMA-1 and Pax-2 (PRB-276P; Covance) antibodies as described by Saarela *et al*. 2017 (Saarela, U., Akram, Desgrange, Rak-Raszewska, Shan, Cereghini, Ronkainen, Heikkilä, Skovorodkin, & Vainio, 2017b).

### Sequence alignment and protein structure prediction

Protein sequences for mouse and human TSP1 and ColXVIII TSP1-like domain (Laminin G-like) were retrieved from Uniprot (The UniProt Consortium, 2019). Sequence alignment was carried out using Clustal Omega (Sievers et al., 2011) with default settings.

A homology model for ColXVIII TSP1 domain was generated with PHYRE2 (Kelley et al., 2015) intensive modeling mode. The mouse ColXVIII TSP1-like domain sequence (The UniProt Consortium, 2019) was submitted for modelling, returning a model based on the crystal structure of the N-terminal domain of collagen IX (PDB code 2UUR). The model had a confidence score of 100 % with 92 % coverage demonstrating an appropriate model. The superimposed image was created using PyMOL (Schrödinger L.L.C. The PyMOL Molecular Graphics System. Schrödinger; New York, NY, USA: 2009. version 1.2r3pre.).

### Statistical analyses

The statistical analyses were performed with the GraphPad Prism 5 program (GraphPad Software, San Diego, California, USA). The Mann-Whitney U test, with eventual Welch’s correction where requested, was used to calculate statistical significances between two groups, and the Kruskall-Wallis test with Dunn’s multiple comparison test was used to compare multiple groups. Values were considered as significant at P-values less than 0.05. In the box plots, whiskers indicate minimum and maximum values, the box borders indicate the lower and upper quarters and the line in the middle of the box the mean value. In the dot plots, whiskers indicate standard deviation (s.d.) and line in the middle the mean value. In the columns, mean is shown with whiskers indicating s.d.

## ACKNOWLEDGEMENTS

The authors wish to thank the Laboratory Animal Centre of the University of Oulu, Biocenter Oulu transgenic core facility, a member of Biocenter Finland and Infrafrontier-EMMA, Biocenter Oulu Light microscopy core facility, a member of Biocenter Finland biological imaging platform services, Biocenter Oulu Virus core facility and Vimma service of the University of Oulu, as well as Aila White, Sirkka Vilmi, Maija Seppänen, Jaana Peters and Päivi Tuomaala for their technical assistance.

## AUTHOR CONTRIBUTIONS

M.M.R-J. and H.J.R. carried out experiments and analysed the data. M.M.R-J., F.N., H.J.R. and T.A.P. designed the study. F.N. performed the N-terminal fragment cultures, part of the cell cycle analyses and dissected and prepared E10.5 AGMs for cultures and E11.5 MMs for qPCR. S.U.A. developed the computer program for counting Six2+ cells. J.T.K. performed the sequence alignment and protein structure modelling. T.S. designed and produced the ColXVIII antibody. I.P. provided OPT analysis support. H.P.E. designed ColXVIII *in situ* probes and N-terminal recombinant proteins. I.V. helped with flow cytometry analysis. V-P.R. helped with microscopy imaging. I.K. produced and purified the N-terminal fragment. A.M. and S.J.V. provided advice on study design. M.M.R-J. made the figures and M.M.R-J., F.N., T.A.P. and H.J.R. wrote the paper. All authors approved the final version of the manuscript.

---

**Figure 2 – figure supplement 1.**
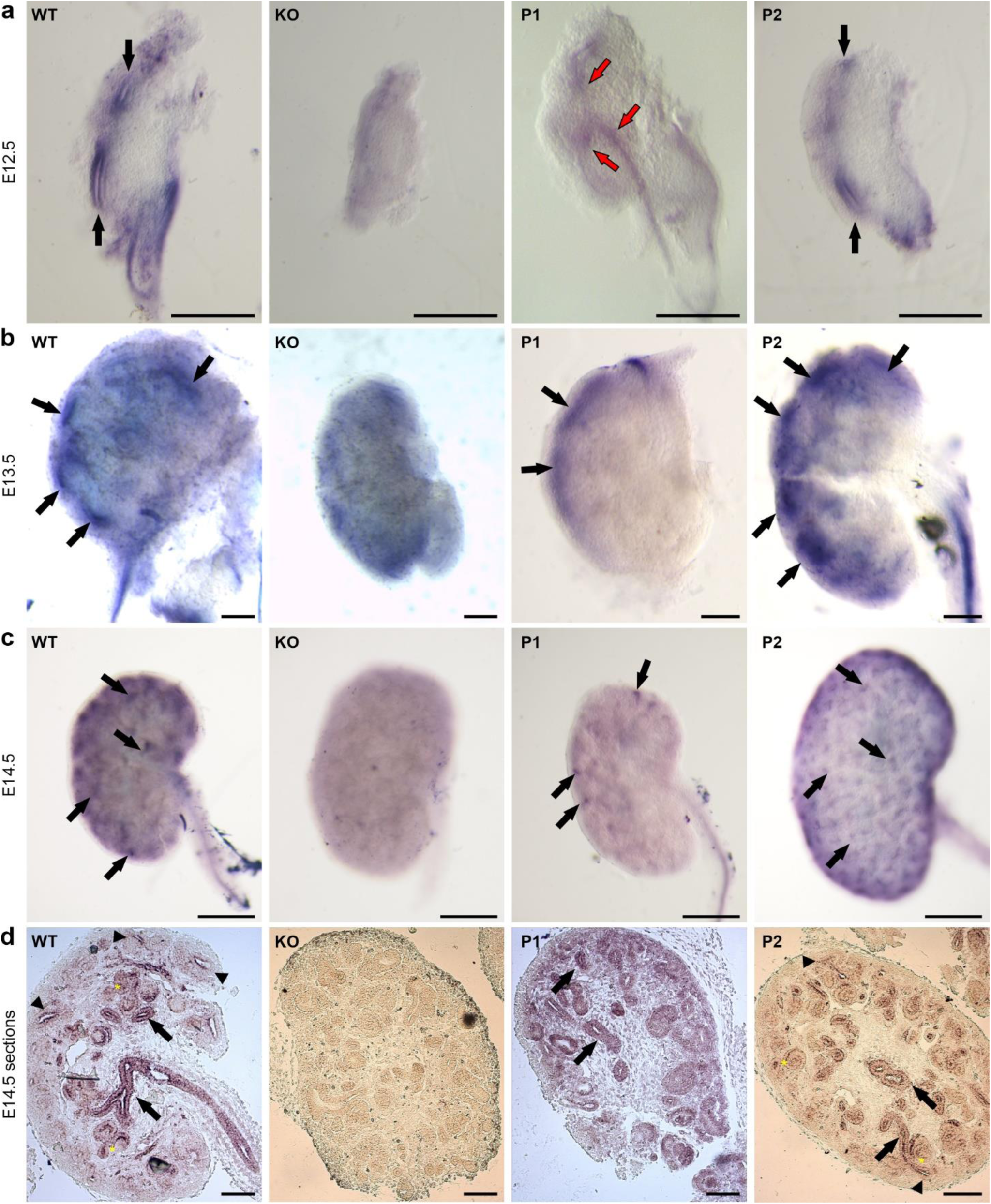
The short ColXVIII form was expressed in the branching ureteric tips. (a) Whole-mount *in situ* hybridisation of E12.5 kidneys with a ColXVIII antisense RNA probe. Black arrows indicate ColXVIII-positive ureteric tips in WT and P2 kidneys. Red arrows indicate ColXVIII-positive stalk area in a P1 kidney, where the ColXVIII signal was not seen in the tips. Bar: 500 μm. (b) Whole-mount *in situ* hybridisation of E13.5 kidneys (bar: 200 μm) and (c) E14.5 kidneys (bar: 500 μm) with the ColXVIII probe. Arrows indicate ColXVIII-positive tubular structures. Note that P1 kidneys had significantly less ColXVIII-positive structures in the surface. (a-c) In the KO kidneys, there was no specific ColXVIII expression. (d) Section *in situ* hybridisation of E14.5 kidneys. In the WT, the ColXVIII expression was seen in the tubules (black arrows), tips (arrowheads) and forming glomerular structures (yellow stars), and the same pattern was seen in the P2 kidneys. In the P1 mice, the ColXVIII expression was mainly seen in the tubules (black arrows). No expression was seen in the KO kidneys. Bar: 100 μm.

**Figure 6 – figure supplement 1.**
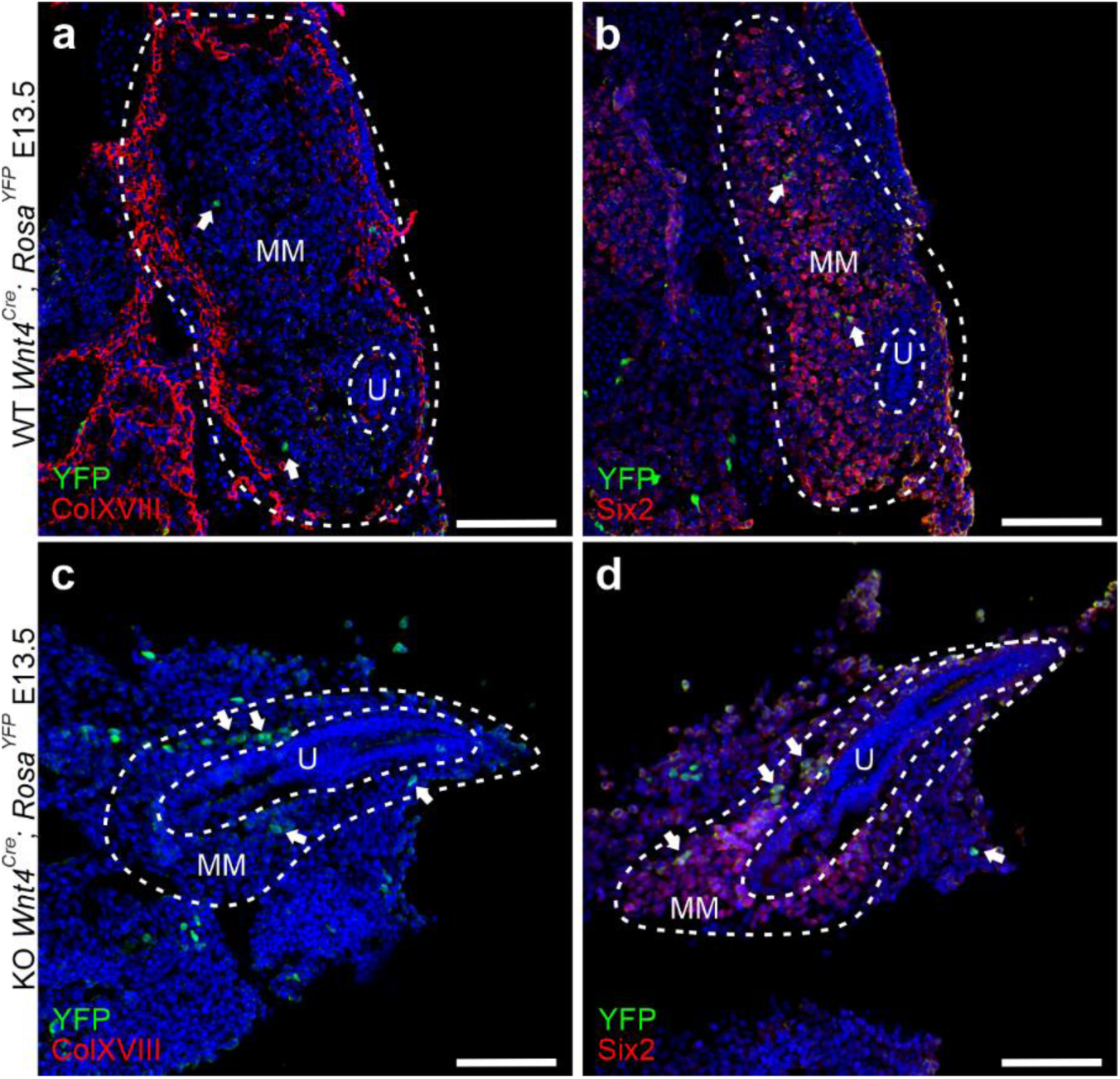
Only a few Wnt4-positive cells were detected at E11.5 in the NPC population. (a) In the WT kidneys, the ColXVIII signal was seen around the ureter (U, dashed line) and in the renal capsule. Only a few Wnt4+-cells (YFP-positive, arrows) were detected in the metanephric mesenchyme (MM, dashed line). (b) Six2+ NPCs were detected throughout the MM, and a few of them were positive for Wnt4 (arrows). (c) No ColXVIII signal was detected in the KO kidneys. Some Wnt4+ cells (arrows) were detected in the MM. (d) Six2+ NPCs were detected in the MM of the KO kidneys and, similarly to the WT, a few of them were positive also for Wnt4 (arrows). Bar = 100 μm

**Figure 7 – figure supplement 1.**
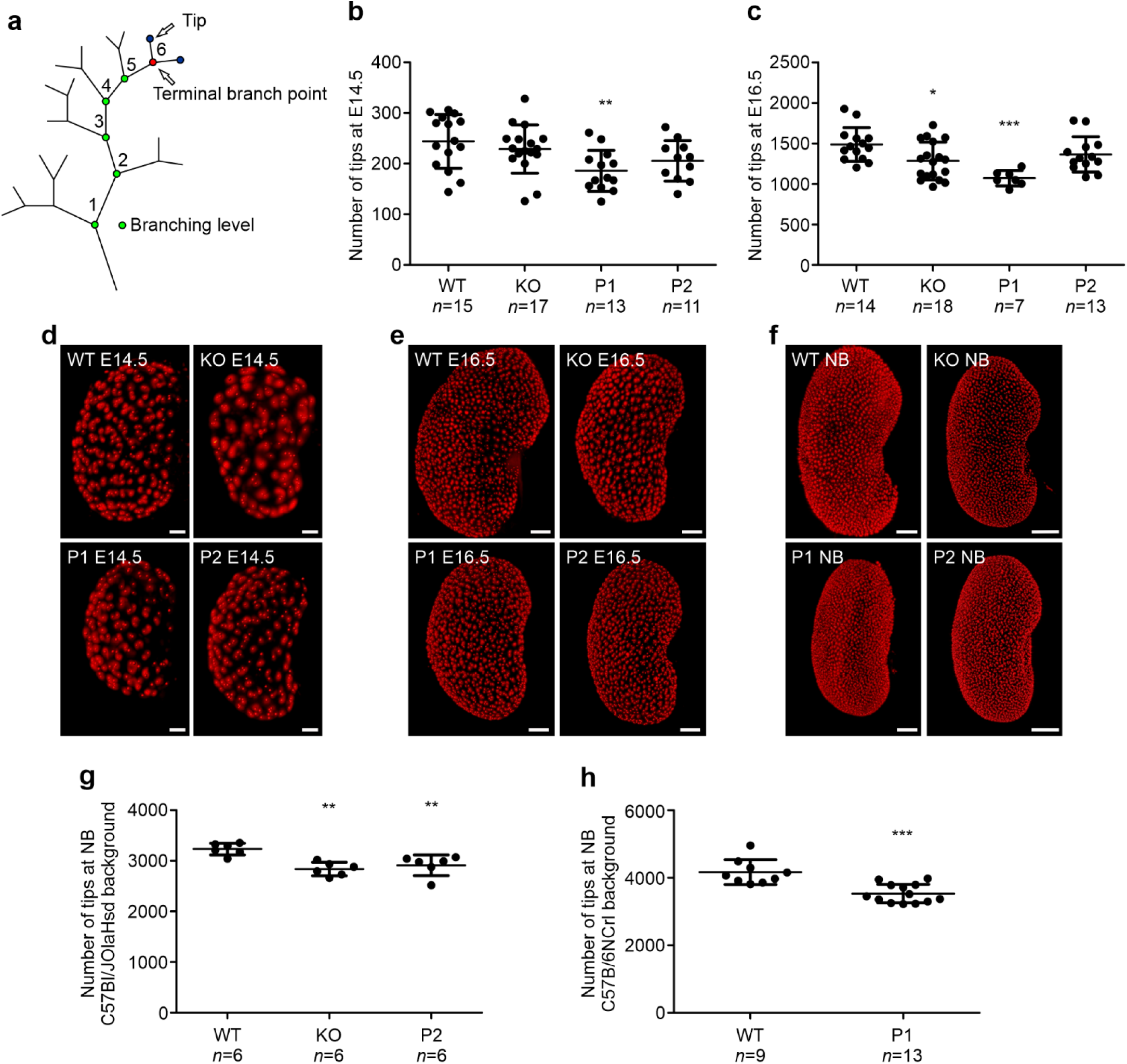
Lack of ColXVIII or its specific isoforms led to reduced branching during the kidney development. (a) A schematic image of analysed parameters from TROMA-1 immunostained and Wnt11 *in situ* hybridised kidneys by OPT. (b) The number of ureteric tips at E14.5 of Wnt11 *in situ* hybridised kidneys analysed by OPT; *n*(WT)=15, *n*(KO)=17, *n*(P1)=13, *n*(P2)=11. (c) The number of Wnt11-positive ureteric tips at E16.5; *n*(WT)=14, *n*(KO)=18, *n*(P1)=7, *n*(P2)=13. (d) Examples of OPT images of Wnt11 *in situ* hybridised (d) E14.5 (bar: 100 µm), (e) E16.5 (bar: 200 µm), and (f) new born (NB) mouse (bar: 400 µm) kidneys used for analysation of the ureteric tip number. (g) The number of ureteric tips in NB WT, KO, and P2 kidneys in a C57Bl/JOlaHsd background. *N*=6 per genotype. (h) The number of ureteric tips of NB WT and P1 kidneys in a C57Bl/6NCrl background. *N*(WT)=9, *n*(P1)=13. The NB graphs are shown separately since the background of the mice was changed during the study, and the kidneys in the C57B/6NCrl line were bigger than those in the C57Bl/JOlaHsd line, but the ratio in the tip number between WT and mutants remained the same. In graphs, means±s.d. are shown. \**P*<0.05, \*\**P*<0.01, and \*\*\**P*<0.001 (Mann-Whitney U – test).

**Figure 7 - figure supplement 2.**
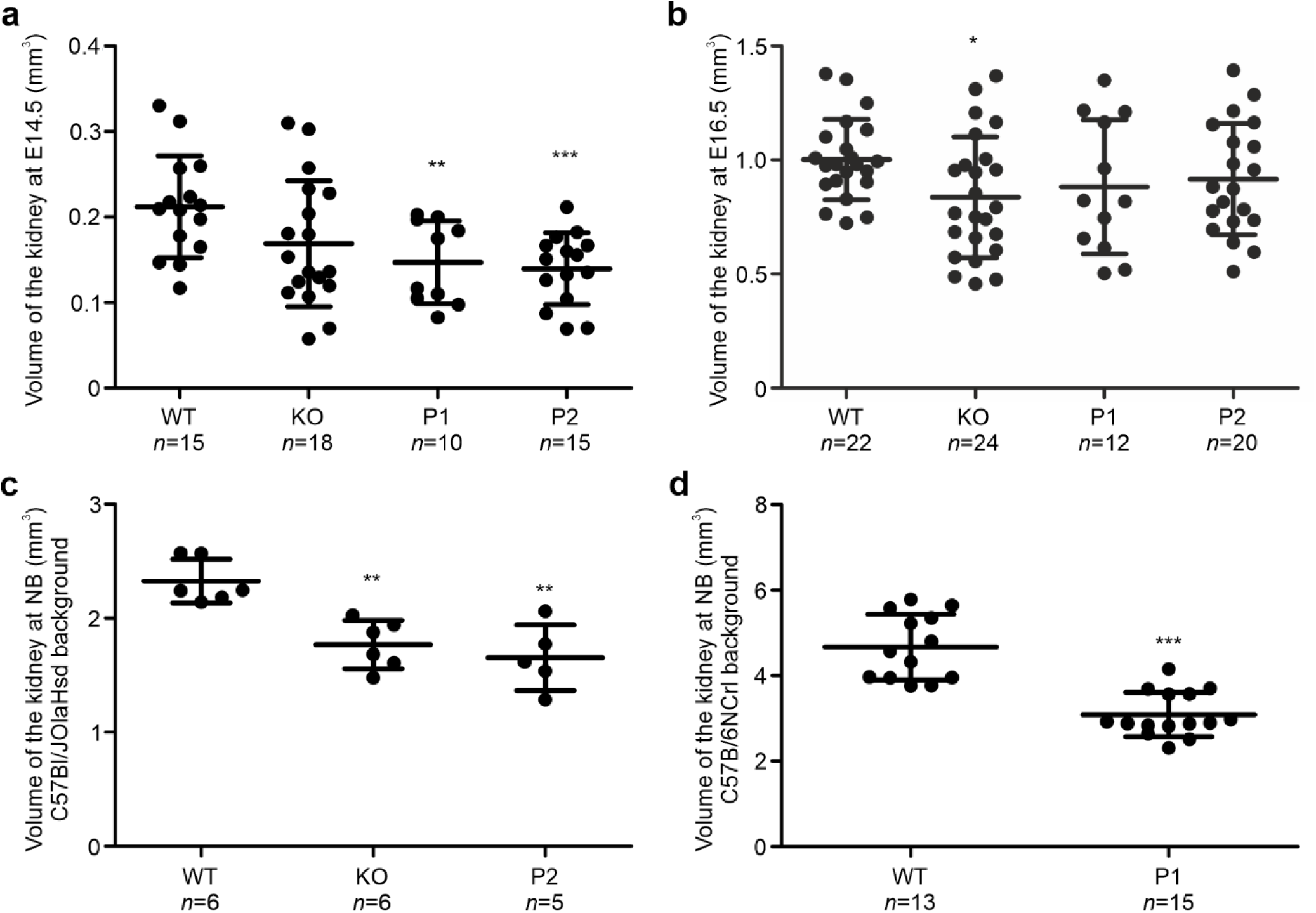
Kidneys without all ColXVIII forms (KO), the short form (P1), or two longer forms (P2) were smaller than WT kidneys throughout the development. The volumes of the kidneys were calculated from Wnt11 *in situ* hybridisation and OPT. WT and ColXVIII mutant kidneys in a C57Bl/JOlaHsd background at (a) E14.5 (*n*(WT)=15, *n*(KO)=18, *n*(P1)=10, *n*(P2)=15), (b) E16.5 (*n*(WT)=22, *n*(KO)=24, *n*(P1)=12, *n*(P2)=20), and (c) NB (*n(WT)*=6, *n*(KO)=6 and *n*(P2)=5. (d) The volume of the NB WT and P1 kidneys in a C57Bl/6NCrl background, *n*(WT)=13, *n*(KO)=15. The NB graphs are shown separately since the kidneys in the C57Bl/6NCrl line were bigger than those in the C57Bl/JOlaHsd line even if the difference between WT and mutants remained the same. In graphs, means±s.d. are shown. \**P*<0.05, \*\**P*<0.01, and \*\*\**P*<0.001 (Mann-Whitney U –test).

**Figure 9 – figure supplement 1.**
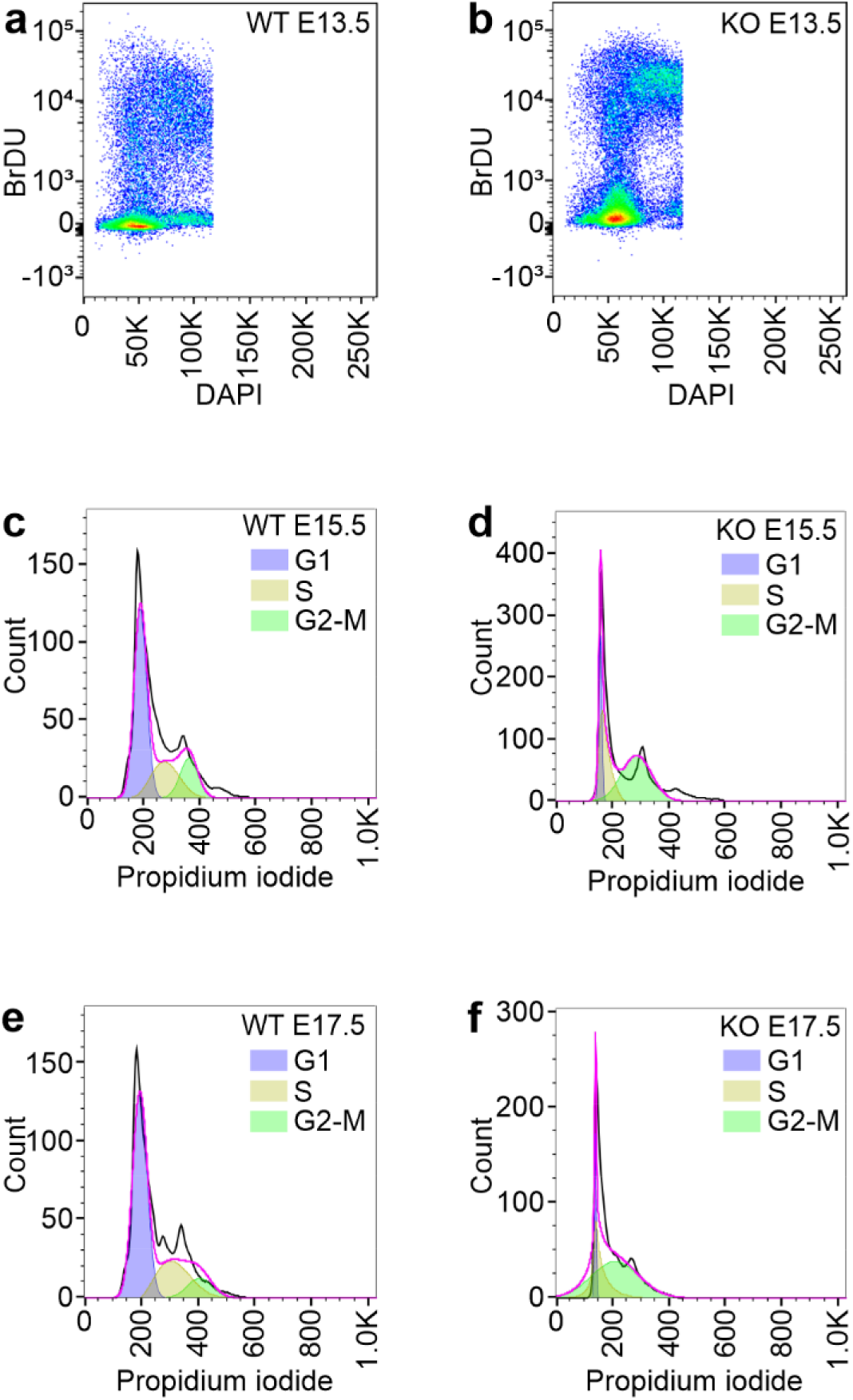
Representative images of cell cycle analyses of Six2-positive cells. Representative images of the Six2+ cell cycle analyses of the E13.5 WT (a) and KO (b) kidneys, E15.5 WT (c) and KO (d) kidneys, and E17.5 WT (e) and KO (f) kidneys. BrDU= 5-bromo-2’-deoxyuridine.

## Movie captions

**Movie 1. A representative time-lapse video of the WT aorta-gonad-mesonephros (AGM) culture.** The AGMs are oriented horisontally, where the limb end is on left. One kidney forms in the middle upper part of the AGM, where the outgrowth of the ureter bud can be seen 9 h onwards. To the lower part forms three kidneys next to each other. The YFP signal is detected in the MM around 10 h timepoint in the upper kidney and in the lower kidneys already within the first hours. The signal increases over time in the MM when the kidney grows. The green YFP signal indicates Wnt4-positive cells. Timelapse was captured with 0.5 h intervals, and the total follow-up time was 39 h.

**Movie 2. A representative time-lapse video of the ColXVIII KO aorta-gonad-mesonephros (AGM) culture.** The AGM is oriented horizontally, where the limbs are located in the right upper part of the imaged area. The kidney is formed on the right lower part of the imaged area in the end of the nephric duct. The YFP signal is detected in the MM of the forming kidney around 17-18 h timepoint and the signal increases in the MM over time when the kidney grows. The green YFP signal indicates Wnt4-positive cells. Time-lapse was captured with 0.5 h intervals, and the total follow-up time was 39 h.

**Movie 3. Example of one nephrogenic niche showing Six2-positive cells in a cap mesenchyme of an E15.5 kidney.** The kidney cortex was imaged through with the multiphoton microscope, and Six2-positive cells in each niche were calculated using a computer program designed for this purpose. The program marks every calculated cell by different colored circles through the stacks and, thus, calculates each cell separately. The cell count comes out in a separate text file in addition to the movie that it produces about the calculation.

